# Transcriptional changes in *Plasmodium falciparum* upon conditional knock down of mitochondrial ribosomal proteins RSM22 and L23

**DOI:** 10.1101/2022.02.09.479728

**Authors:** Swati Dass, Michael W. Mather, Liqin Ling, Akhil B. Vaidya, Hangjun Ke

## Abstract

The mitochondrion of malaria parasites is an attractive antimalarial drug target, which require mitoribosomes to translate genes encoded in the mitochondrial (mt) DNA. *Plasmodium* mitoribosomes are composed of highly fragmented ribosomal RNA (rRNA) encoded in the mtDNA. All mitoribosomal proteins (MRPs) and other assembly factors are encoded in the nuclear genome. Here, we have studied one small subunit (SSU) MRP, RSM22 (Pf3D7_1027200) and one large subunit (LSU) MRP, L23 (Pf3D7_1239100) in *Plasmodium falciparum*. We show that both proteins localize to the mitochondrion and are essential for parasite survival. However, the parasites survive conditional knock down (KD) of PfRSM22 for two days longer than PfMRPL23 KD. RNA sequencing revealed transcriptomic changes of the nuclear and mitochondrial genomes upon KD of these MRPs. In the early phase following KD, while most mt rRNAs and transcripts of putative MRPs were downregulated in the absence of PfRSM22, several mt rRNAs and MRPs were upregulated after KD of PfMRPL23. At the late time points of KD, loss of PfRSM22 and PfMRPL23 caused defects in many essential metabolic pathways, leading to parasite death. There was a significant overlap among the mitochondrial related transcripts downregulated in the late phase of PfRSM22 and PfMRPL23 KDs. We have also identified a list of mitochondrial proteins of unknown function that are likely *Plasmodium* MRPs based on their structural similarity to known MRPs as well as their expression profiles in KD parasites.

## INTRODUCTION

The mitochondrion is an essential organelle in eukaryotes that emerged from an alpha proteobacterial endosymbiont ~ two billion years ago (1). It has diversified tremendously with every evolving eukaryote adapting the mitochondrion’s roles according to the physiological demands of each organism’s evolutionary niche (2). A vast literature is available on understanding the composition and physiology of mitochondria in model eukaryotic systems (3,4). This information has been exploited to help understand the evolutionary and functional differences in the mitochondria of less studied eukaryotes, including *Plasmodium spp.,*the causative agent of malaria, but much remains unknown about the function of organelles in these early diverging organisms (5,6).

According to the latest World Malaria Report, there has been a spike of 14 million malaria cases in 2020, raising the death toll to 627,000 around the world (7). This signifies malaria as a heightened global health burden and demands a thorough understanding of *Plasmodium* biology to support advances in antimalarial drug development. The *Plasmodium* mitochondrion has been validated as an antimalarial drug target due to its essentiality in the parasite and differences from the human counterpart. The parasite mitochondrial electron transport chain (mtETC) is a target of an array of clinical and pre-clinical antimalarials, such as atovaquone and ELQ-300, which do not affect the human mtETC at pharmacologically relevant concentrations (8,9). Despite its importance, our understanding of the *Plasmodium* mitochondrion remains limited (10). The *Plasmodium* mitochondrial genome encodes only three mtETC proteins (*cyt b,* COX I, and COX III) (11). Other than these three proteins, this genome encodes mt rRNA on both strands of the mtDNA (12,13). Unlike the rRNAs of most other organisms, the *Plasmodium* mt rRNAs are fragmented into 39 distinct molecules ranging in length from 20 to 200 bases. Another unusual phenomenon of *Plasmodium* mt rRNA is the presence of a short oligo (A) tail at the 3’ end of most mt rRNA transcripts (14). On the other hand, all MRPs are encoded in the parasite nuclear genome and post translationally transported to the parasite mitochondrion where they assemble along with mt rRNAs to form a functional mitoribosome (15).

Currently, 43 putative MRPs in *Plasmodium falciparum* (Pf) have been annotated based on their sequence similarity with known MRPs of bacterial and mitochondrial origin (www.PlasmoDB.org). However, the species-specific *Plasmodium* MRPs remain entirely unknown (16,17). We have previously studied four annotated MRPs in *P. falciparum*, including PfMRPL13, PfMRPS12, PfMRPS17, and PfMRPS 18 (18,19). Our data show that knocking down PfMRPs reduces the cytochrome *c* reductase activity of the *bc_1_* complex, consistent with an essential role of mitoribosomes in translating mtETC proteins (18). We have also shown that disrupting *P. falciparum* mitoribosomes impairs mitochondrial metabolic pathways, rendering the parasite hypersensitive to antimalarial drugs targeting the mtETC (19). However, it remains unknown how the loss of mitoribosomes affects the overall health of the parasite, gradually leading to parasite demise. Apart from the four MRPs mentioned above, the other 39 MRPs have not been characterized in *Plasmodium*.

Here, we have studied two annotated MRPs, PfRSM22 and PfMRPL23. We used CRISPR/Cas9 and the TetR-DOZI-aptamer system (20) to generate conditional KD lines to control expression of PfRSM22 and PfMRPL23 and insert a triple hemagglutinin (3xHA) tag at the C terminal end of the proteins. Using these transgenic parasite lines, we confirm that PfRSM22 and PfMRPL23 are localized to the mitochondrion and are essential for parasite development and survival. To understand the global and mitochondrial transcriptomic changes upon KD of PfRSM22 or PfMRPL23, we performed RNA sequencing of the asexual stage parasites at multiple time points. We have categorized the parasite morphological and transcriptomic changes at the early and late phases of KD. In addition, we identified differentially regulated *Plasmodium* proteins of unknown function that exhibited structural similarity to distant MRPs of other organisms. These proteins could represent new *Plasmodium* MRP candidates.

## RESULTS AND DISCUSSION

### PfRSMS22 and PfMRPL23 are essential mitochondrial proteins

PfRSMS22 (Pf3D7_1027200) is annotated as a putative MRP S22 in *Plasmodium* database (www.PlasmoDB.org), but contains an S-adenosyl-L-methionine (SAM) dependent methyltransferase like domain (InterPro IPR029063) (21). Its ortholog in *Homo sapiens* is annotated as methyltransferase like protein 17 (METTL17); however, orthologues of PfRSM22 found in yeast, *Tetrahymena, Trypanosoma* and *Toxoplasma* are annotated as RSM22 (22–24). The methyltransferase activity of this protein is known to play a role in maturation of mt rRNA and mitoribosomal assembly (21). *Trypanosoma brucei* RSM22 interacts with immature rRNA and is a member of the early SSU assemblosome. Its KD leads to reduction of the 9S rRNA (SSU mt rRNA) expression level. However, its association with the mature mitoribosome is not observed by cryo-EM (24). We propose that Pf3D7_1027200, although annotated as putative MRPS22 in PlasmoDB, should be termed RSM22, a putative mitochondrial ribosome associated protein on the basis of its resemblance to the InterPro Ribosomal protein Rsm22-like protein family (IPR015324/PF09243) with strong statistical support (E = 0.0) (Figure S1) (25). mS22, found in metazoan mitoribosomes, belongs to a different protein family that lacks methyltransferase activity (InterPro IPR019374).

PfMRPL23 (Pf3D7_1239100) likely belongs to LSU based on its homology to L23 proteins, which are conserved throughout all kingdoms of life (InterPro Ribosomal protein uL23 family IPR013025/Pfam PF00276, E=1.1e-13 for the PfMRPL23 match) (Figure S2). Previous reports represent that *E. coli* L23 interacts with domain III of the LSU rRNA and is a part of the exit tunnel of the ribosome (26,27). The exit tunnel is a crucial site for the mitoribosome as it is involved in co-translational insertion of highly hydrophobic mtETC proteins into the mitochondrial inner membrane (28).

To investigate the essentiality of PfRSMS22 and PfMRPL23, we genetically modified their loci in the *P. falciparum* D10 WT parasite line using a CRISPR/Cas9 mediated double crossover recombination strategy (20). PfRSMS22 and PfMRPL23 loci were individually modified by integrating 3xHA and 8 copies of TetR binding RNA aptamers at the 3’ end of the coding sequence, providing controlled expression of the gene using the TetR-DOZI-aptamer system (29). Positively transfected parasites were obtained in 4 weeks post transfection under constant pressure of blasticidin (BSD) and anhydrotetracycline (aTc, 250 nM). Genotyping of the transgenic lines, D10-PfRSM22-3HA and D10-PfMRPL23-3HA, was performed using specific primers (Table S1). The strategy of integration, confirmation of modified loci and the absence of the wildtype gene are shown in Figure S3A and S3B. Intracellular localization of PfRSM22-3HA and PfMRPL23-3HA was examined by immunofluorescence assay (IFA). Both proteins co-localized with the fluorescent mitochondrial probe Mitotracker red (Figure 1A and 1B), indicating that PfRSM22 and PfMRPL23 are mitochondrial proteins.

**Figure 1.**
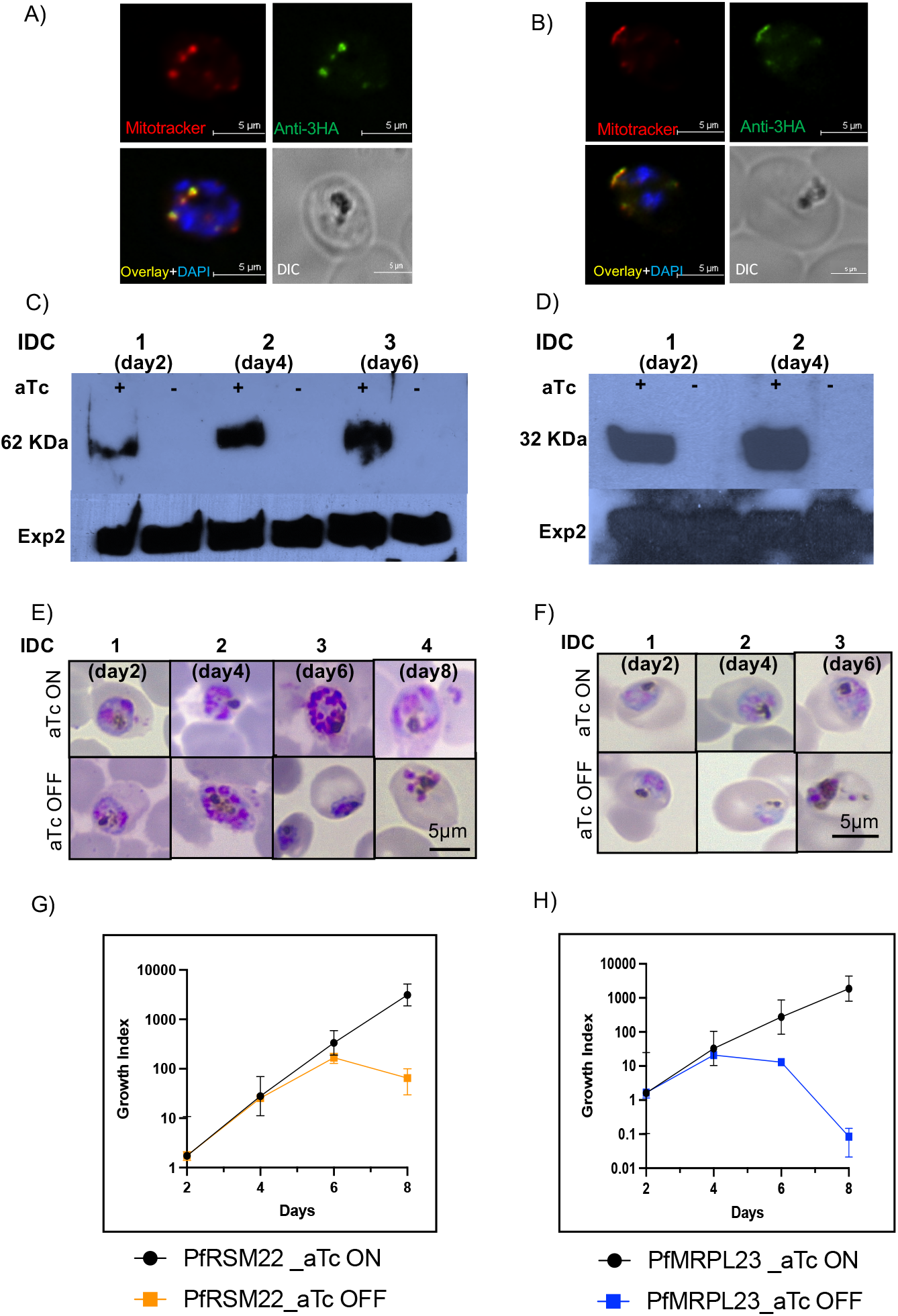
PfRSMS22 and PfMRPL23 are localized to the mitochondrion and are essential for the survival of the parasite.

Upon removal of aTc from the culture medium to induce KD of PfRSM22 or PfMRPL23, we assessed parasite viability and morphology over 8 days, or 4 intraerythrocytic cycles (IDCs). The parasites were grown in medium containing 250 nM aTc to allow expression of the respective proteins (aTc ON). As shown in Figure 1C and 1D, PfRSM22 and PfMRPL23 were expressed in presence of aTc, and within 2 days of aTc removal, neither proteins could be detected by Western blotting. The effects on parasite morphology observed over time following KD of PfRSM22 and PfMRPL23 are shown in Figure 1E and 1F using light microscopy of Giemsa-stained thin blood smears. Late-stage parasites grown without aTc were smaller and morphologically abnormal compared to aTc plus parasites by day 6 in the D 10-PfRSM22-3HA line and by day 4 in the D10-PfMRPL23-3HA line. It was interesting to note the difference in progression to death upon KD of PfRSM22 vs PfMRPL23. We found that D10-PfRSM22-3HA survived for 6 days and appeared dead on the 8^th^ day after aTc removal, while D10-PfMRPL23-3HA parasites died after day 4 following aTc removal. Overall, we show that both PfRSM22 and PfMRPL23 are essential for the survival and development of asexual stage parasites. However, PfRSM22 KD parasites were able to survive for one IDC more than the PfMRPL23 KD parasites. Quantification of parasitemia in both KD lines is shown in Figure 1G and 1H.

Based on these results, we have categorized the effects of knocking down PfRSM22 or PfMRPL23 into two categories, the early phase (day 2 aTc off) and the late phase (day 6 aTc off for PfRSM22 vs day 4 aTc off for PfMRPL23). In both parasite lines, morphological and growth defects became evident only in the late phase after KD. In the early phase after KD, however, both KD parasite lines appeared healthy despite PfRSM22 and PfMRPL23 proteins being undetected by Western blotting. We reasoned that the effects of mitoribosomal synthesis inhibition are probably not lethal until the preexisting protein translation products (mtETC proteins) have been entirely lost to protein turnover. Previous studies have shown that mitochondrial respiration occurs at a low level in asexual blood stages (30) and functions mainly to serve pyrimidine biosynthesis by recycling reduced ubiquinone (31,32). Thus, continued inheritance of previously assembled mtETC complexes can serve this function for at least one generation without the need for newly assembled mitoribosomes.

### Transcriptomic changes upon KD of PfRSM22 or PfMRPL23 in the early phase

To investigate global transcriptional changes upon loss of essential mitoribosomal proteins, we tightly synchronized both the D10-PfRSM22-3HA and D10-PfMRPL23-3HA lines and initiated KD by removing aTc from trophozoite stage parasites (day 0). We performed RNA sequencing of total polyA+ RNA isolated from PfRSM22 KD parasite on days 2, 4 and 6, and PfMRPL23 KD parasites on days 2 and 4 (Experimental Procedures). RNA samples isolated from D10-PfRSM22-3HA and D10-PfMRPL23-3HA parasite lines grown in the presence of aTc for two days were used as controls for RNA sequencing. Corresponding to the Western blot results in Figure 1C and 1D, PfRSM22 and PfMRPL23 transcripts were downregulated on all days following KD (Figure S4). Normalized read counts and DEseq2 values of the RNA sequencing results are provided in Table S2. Differentially regulated genes with statistical significance (Benjamini-Hochberg p-adjusted value) below 0.05 were selected to generate Volcano plots (Table S2).

Volcano plots represent statistical significance (p adj) vs the magnitude of changes (log2 fold change) of all gene transcripts in PfRSM22 aTc OFF vs PfRSM22 aTc ON samples and PfMRPL23 aTc OFF vs PfMRPL23 aTc ON samples. Each dot represents one gene in the Volcano plots. On day 2 post aTc removal, we observed significant transcriptomic alterations in both KD parasite lines. Figure 2A depicts that, within 2 days of PfRSM22 KD, 1257 genes were significantly upregulated, and 1239 genes were significantly downregulated. The magnitude of the changes in the expression levels ≥ 2-fold (136 genes) and ≤ 2-fold (581 genes) are shown as black dots. Likewise, within 2 days of PfMRPL23 KD, Volcano plot shows that a total of 1612 genes significantly upregulated and 1591 genes significantly downregulated (Figure 2B). About half of these differentially regulated transcripts were log2 fold ≥ 1 or log2 fold ≤-1 (Figure 2B). We observed that 23 downregulated genes belong to the broader KEGG pathway representing proteasome catabolic processes of cellular proteins (Figure 2C). Furthermore, structural constituents of ribosomes, RNA binding proteins and translation related proteins (80 genes in total) were also downregulated; these proteins belong to the ribosome KEGG pathway (Figure 2C, orange). The PfMRPL23 KD downregulated a wider range of pathways on day 2. KEGG analysis revealed that transcripts related to the pentose phosphate pathway, RNA degradation and spliceosome proteins were significantly downregulated, along with the ribosomal biogenesis pathway (Figure 2C, blue).

**Figure 2.**
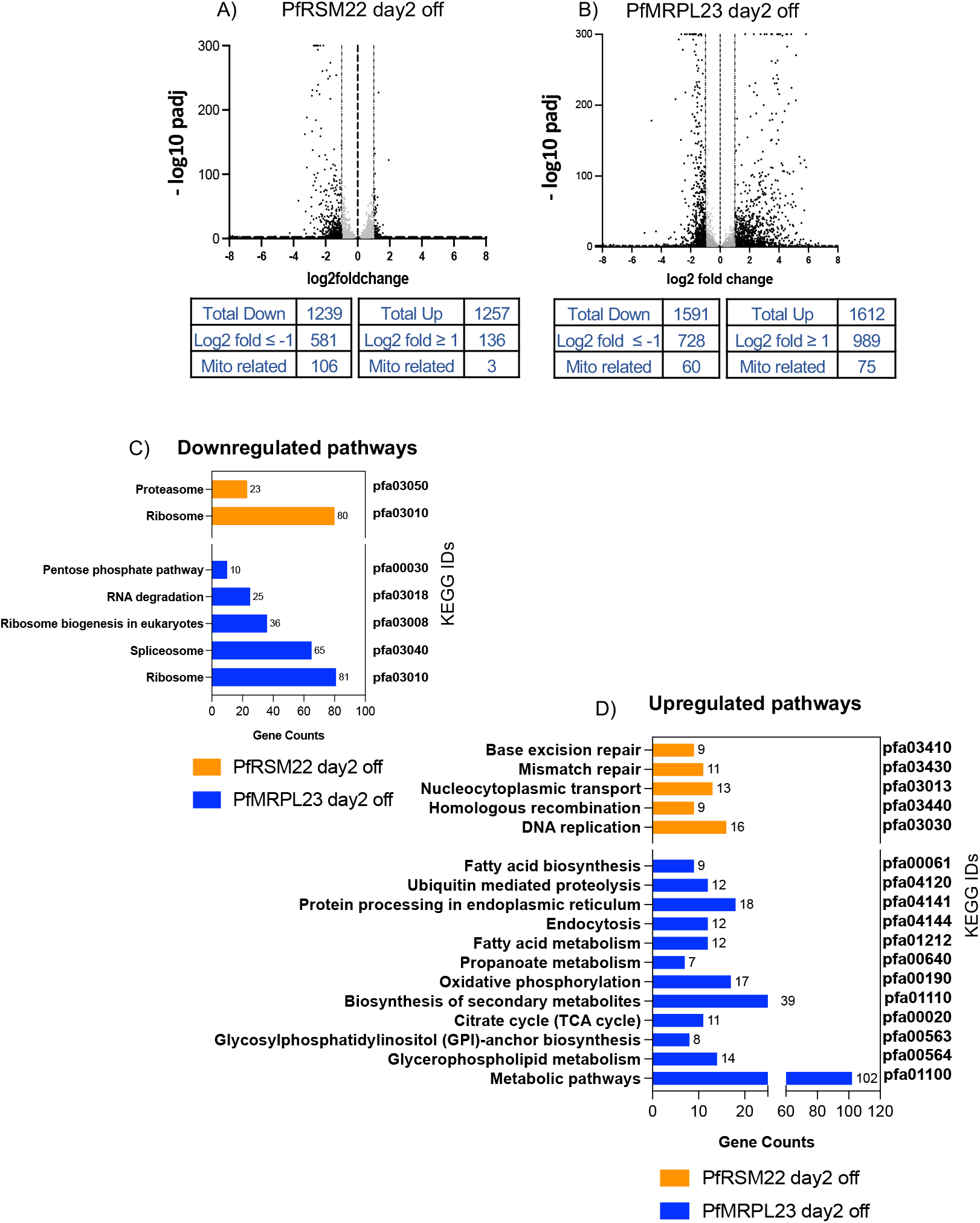
Transcriptomic changes in the early phase after KD of PfRSM22 and PfMRPL23.

Among the upregulated pathways, on the other hand, the parasites showed disparity between PfRSM22 and PfMRPL23 KD at the transcript level. The PfRSM22 KD on day 2 resulted in a slight upregulation of transcripts involved in DNA metabolic processes and nucleocytoplasmic transport, suggesting increase in DNA metabolic processes (Figure 2D, orange). On the other hand, the PfMRPL23 day2 KD showed an increase in multiple metabolic processes, such as fatty acid metabolism, TCA cycle, protein processing in ER, oxidative glycerophospholipid metabolism, biosynthesis of secondary metabolites and other metabolic pathways (Figure 2D, blue).

Overall, these data suggest that KD of PfRSM22 and PfMRPL23 leads to dramatic changes at the transcriptional level as early as day 2, when minimal morphological changes were noticeable, and parasites appeared healthy (Figure 1 E and F). Further work would be required to correlate the transcriptional changes observed in biological pathways to alterations at the protein levels.

### Early effects of PfRSM22 and PfMRPL23 KD on transcripts of mitoribosomal components

There are 43 annotated *Plasmodium* MRPs, of which 14 are SSU proteins and 29 are LSU proteins, based on homology with bacterial ribosomes and mitoribosomes of other eukaryotes. This is close to half the number of proteins known in yeast and mammalian mitoribosomes (33). The MRPs have been categorized into early, intermediate and late assembly proteins based on their hierarchical association at different time points forming multiple pre-ribosomal complexes (33,34). While nothing is known about the mitoribosomal assembly intermediates in *Plasmodium*, previous research in other organisms describe the hierarchy of mitoribosomal assembly elaborately (34–36). RSM22 is known to play a role in early assembly and maturation of mt SSU (24). L23 is one of the intermediate binding proteins that makes few or no contact with early assembly proteins, but their incorporation into the ribosome depends on stability of early assembly proteins (37).

We found that 24 out of 43 annotated PfMRPs were significantly downregulated on day 2 KD of PfRSM22; among these, 10 belonged to the SSU and 14 belonged to the LSU (Figure 3A, orange). No PfMRP transcript was upregulated upon PfRSM22 KD. PfMRPL23 KD led to downregulation of transcripts encoding 19 PfMRPs (6 of SSU and 13 of LSU) and upregulation of 5 PfMRPs of which 2 belonged to the SSU (S8 and S14) and 3 were LSU PfMRPs (L11, L16 and L19) (Figure 3A, blue). These data suggest that, in the early phase, PfRSM22 KD resulted in downregulation of most PfMRPs, whereas PfMRPL23 KD caused upregulation of some PfMRPs. The reasons behind potential stabilization or overexpression of mRNA encoding some MRPs while downregulating mRNA for other MRPs remained unexplained at present.

**Figure 3.**
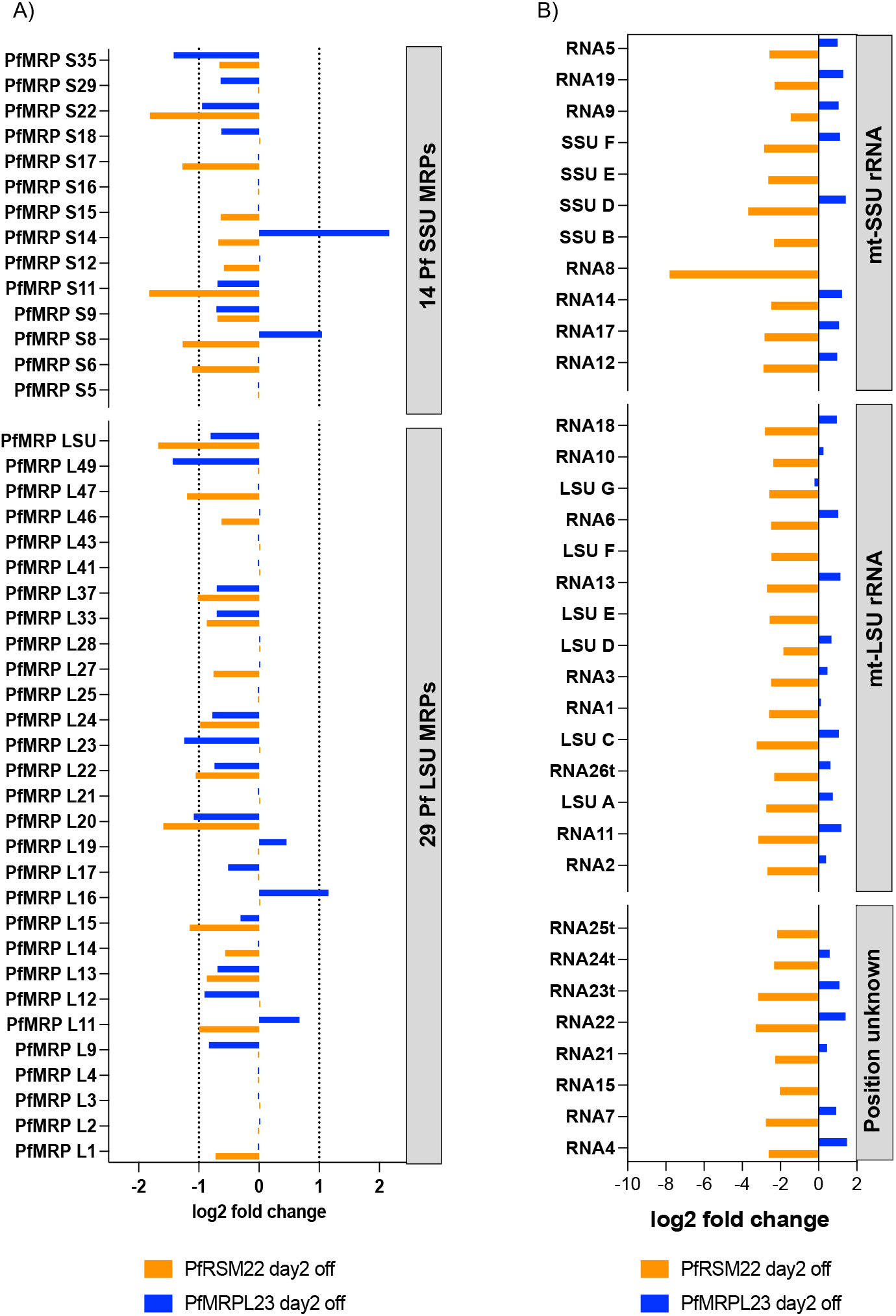
Early effects of PfRSM22 and PfMRPL23 KD on transcripts of mitoribosomal components.

A trend observed in the evolution of mitoribosomes is the reduced size of mt rRNA with an increase in the number of MRPs, as shown in resolved structure of mitoribosomes (38). An illustrative example of this trend is observed in *Trypanosoma brucei*, where the mt rRNAs are as small as 9S (611 nt) and 12S (1150 nt) and associate with more than 120 MRPs to form the mitoribosome (39). Many of these MRPs are found only in those species related to *Trypanosoma*. *Plasmodium* has extended this trend of mt rRNA reduction to an extreme degree, consisting of highly fragmented rRNA pieces (40).

A higher sequencing depth and presence of a poly A tail in most of the mt rRNA fragments facilitated investigation of these transcripts in this study. The PfRSM22 KD resulted in decreased levels of 34 mtDNA encoded rRNA fragments (Figure 3B, orange), which suggests disruption of mitoribosomal components by day 2 post KD. Five mt rRNA fragments (RNA16, RNA20, SSUA, RNA27t, LSUB) could not be detected in our RNA sequencing results. These data were consistent with the fact that 24 PfMRP transcripts were also downregulated at this time point. The result of PfRSM22 KD parallels those of previous studies involving RSM22 disruption in mammalian and *Trypanosoma* which resulted in reduction of SSU mt rRNA levels (24,37). On the other hand, PfMRPL23 KD led to upregulation of 24 mt rRNA transcripts and no significant downregulation of any mt rRNA (Figure 3B, blue), implying that PfMRPL23 KD caused slight stabilization of many mt rRNA transcripts. These data suggested that PfMRPL23 might not be involved in the mitoribosomal assembly process until later time points as suggested in human mitoribosomes (36). The positions of all mt rRNAs on the ribosomal secondary structure proposed in previous studies (12) are shown in Figure S5.

Our RNA sequencing data relating to the mt rRNA fragments further confirm some intriguing aspects of mtDNA transcription in malaria parasites that were noted previously (13,14). Individual mt rRNA fragments do not appear to be expressed in stoichiometric levels, suggesting differential expression profiles of these transcripts as well as their contributions to mitoribosomal assembly in *Plasmodium*. Furthermore, 34 out of 39 fragmented rRNA molecules were sequenced as being poly A+, which was largely in agreement with the previous results of Feagin and colleagues (14) showing the presence of post transcriptional polyadenylation. Unlike transcripts of the apicoplast genome, mt rRNA are not only fragmented but also post-transcriptionally polyadenylated. The exact function and enzymes participating in the process of polyadenylation of mt rRNAs are not known. Our study reiterates the importance of these unanswered questions.

Additionally, we checked the transcriptional changes in the genes related to the parasite apicoplast. We found that four nuclearly encoded apicoplast ribosomal proteins (PfARPs; S14, L12, L18 and L35) and several transcripts encoding proteins targeted to the apicoplast (Figure S6A) were also downregulated in the early phase of PfRSM22 and PfMRPL23 KD. Some PfARPs are, however, encoded in the apicoplast genome, and due to the absence of polyadenylation in apicoplast transcripts, the effect of PfRSM22 and PfMRPL23 KD on apicoplast encoded transcripts remained unexamined here.

Together, in the early phase of PfRSM22 and PfMRPL23 KD, transcripts of mitoribosomal components were significantly perturbed. The findings from this study are limited due to the current incomplete picture of MRPs in *Plasmodium*. These results do not account for unknown MRPs that undoubtedly remain undiscovered in apicomplexan parasites. In addition, PfMRPL23, like a number of other PfMRPs, contains additional stretches of amino acid sequence not present in orthologues from unrelated organisms (Figure S3) (25). The role of parasite specific regions in these proteins, and their interactions with other proteins as well as fragmented mt rRNA remain unexplored at this point.

### Transcriptomic changes upon PfRSM22 and PfMRPL23 KD in the late phase

To understand the transcriptomic changes in the KD parasites right before the onset of parasite death, we carried out RNA sequencing of PfRSM22 KD parasites sampled at day 6 and PfMRPL23 at day 4 following KD. We chose to compare these late phase time points after KD of the respective lines based on similarities in growth decline and abnormal morphology. The transcriptome of PfRSM22 at day 4 after KD was also examined (included in Table S3) but it is not included in the comparison here since there were only a few transcripts that altered compared to data of the day 2 after KD. The overall profile of transcripts upregulated and downregulated upon PfRSM22 KD on day 6 and PfMRPL23 KD day 4 are shown in the Volcano plots in Figure 4A and 4B. Two downregulated KEGG pathways in common to both KDs at the later time points were related to the proteosome and oxidative phosphorylation (OxPhos) (Figure 4C). Downregulation of proteosome related functions suggests defective protein homeostasis at the late phase of KD. Reduction of transcripts related to OxPhos pathway likely to be a downstream effect of collapsing mtETC components due to KD of PfRSMS22 and PfMRPL23 compromising mitochondrial protein synthesis. The GO term pathway related to oxidoreductase activity (GO: 0016491) was significantly downregulated in PfRSM22 and PfMRPL23 KD samples (Table S3). These indicate a series of events emanating from dysregulation of the mitochondrion in the parasite. These observations are aligned with our previous studies that have shown severe defects in cytochrome *bc1* complex activity coinciding with noticeable morphological changes in the parasite due to KD of essential PfMRPs (18,19). Interestingly, PfMRPL23 KD on day 4 also caused downregulation of transcripts involved in metabolic pathways, biosynthesis of secondary metabolites, fatty acid metabolism, GPI anchor biosynthesis, protein processing in ER and biosynthesis of amino acids (Figure 4C, Blue). These findings are consistent with parasites being on the cusp of demise as their metabolism is throttled in response to mitochondrial disruption. Figure 4D represents the KEGG pathways upregulated after PfRSM22 KD on day 6 and after PfMRPL23 KD on day 4, which include transcripts related to cytoplasmic ribosome biogenesis. Specific upregulated KEGG pathways due to PfMRPL23 KD on day 4 included transcripts related to spliceosome and phagosome.

**Figure 4.**
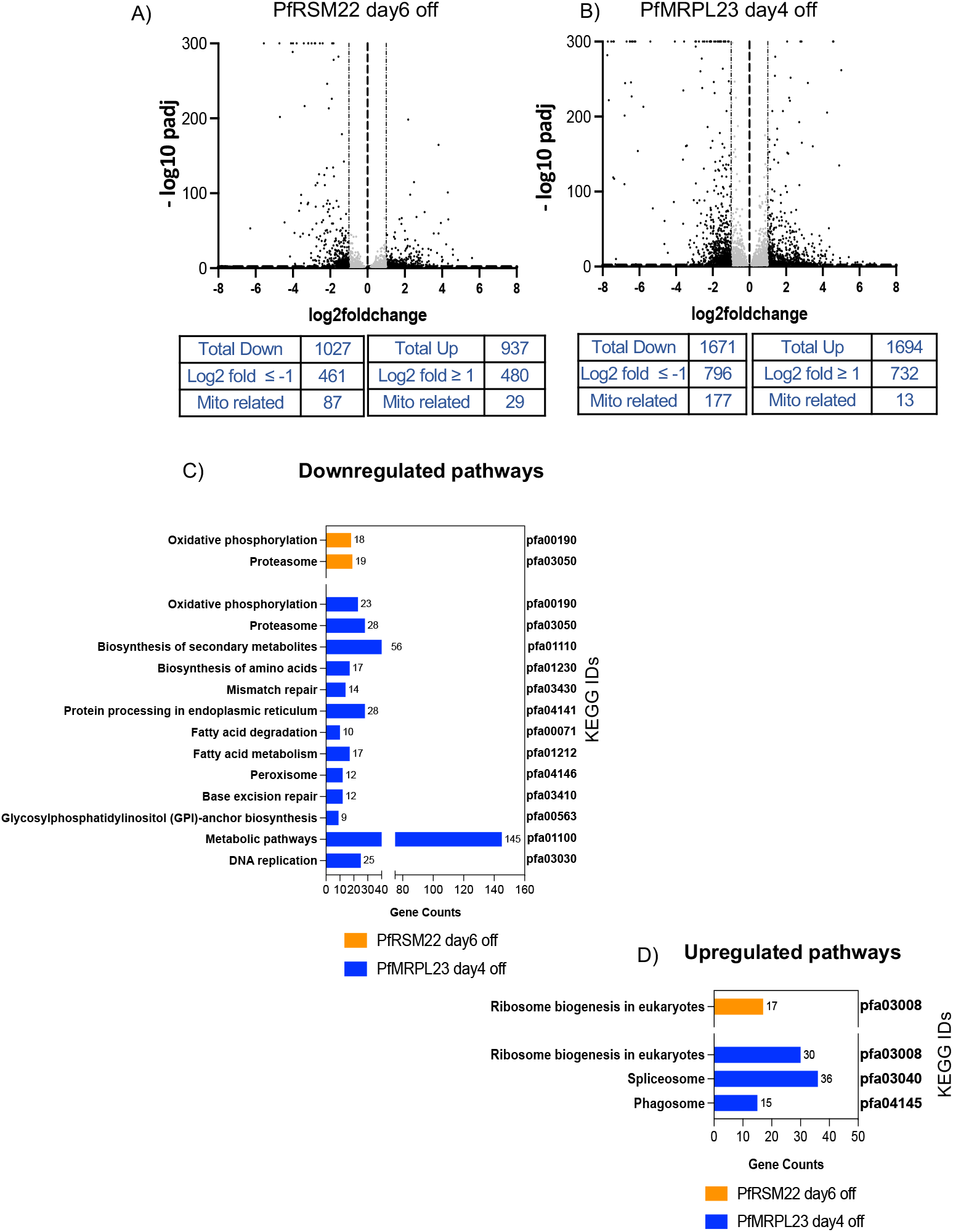
Transcriptomic changes in the late phase after KD of PfRSM22 and PfMRPL23.

We further looked into the effects of the late phase KD on transcripts related to the parasite mitochondrion with the help of current information on the *P. falciparum* mitochondrial proteome. Although the complete mito-proteome of *P. falciparum* remains to be established, recent investigations of organellar proteins based on fractionation and bioinformatic approaches have generated lists of potential nuclearly-encoded proteins that are likely translocated to the parasite mitochondrion. We selected 475 proteins as a putative (unavoidably incomplete) mito-proteome based on manual annotation (41), *Plasmodium* orthologues of *Toxoplasma* mito-proteome (42,43) and selected proteins listed in PlasmoMitoCarta (17). This curated list along with 42 mtDNA encoded transcripts (Table S4) is denoted “mitochondria related transcripts” herein. The list of 517 transcripts was used to assess the mitochondrial components of the transcripts significantly altered in the late phase following both KDs. Figure 5A shows Venn diagrams of significantly downregulated transcripts (log2 fold ≤ −1) observed on day 6 following PfRSM22 KD, on day 4 following PfMRPL23 KD, and in the mitochondria (Table S4). The KD of PfMRPL23 led to highly significant downregulation of mRNAs encoding mitochondrial proteins as well as transcripts encoded on the mtDNA. There were 186 common genes downregulated more than 2-fold on day 6 of the PfRSM22 KD and day 4 of the PfMRPL23 KD.

**Figure 5.**
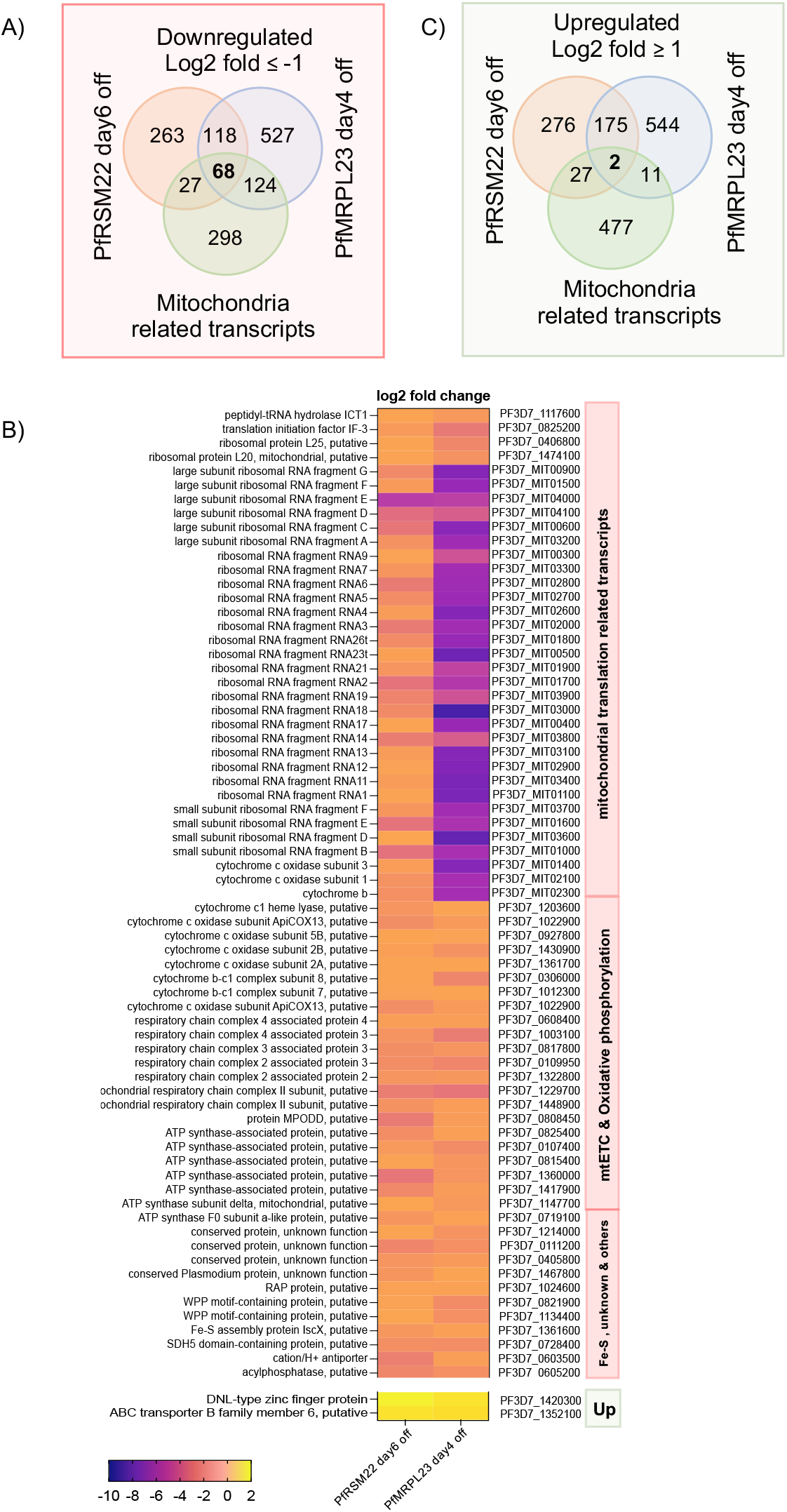
Genes regulated in common after KD of PfRSM22 and PfMRPL23 in the late phase.

Sixty-eight of those 186 commonly downregulated transcripts are related to mitochondria, including mt rRNA transcripts, the peptide release factor PfICT1 (44), translation initiation factor IF-3, PfMRPL20, PfMRPL25, etc. Figure 5B is a heat map representing log2 fold change (log2 fold ≤ −1) in the expression levels of these 68 mitochondria related transcripts. Together, this list suggests defects in mitochondrial protein translation, leading to the downregulation of all three mtDNA encoded transcripts of mtETC. The other mtETC and OxPhos transcripts listed in the heatmap are encoded in nuclear genome which together suggests a gradual collapse of transcripts essential for the parasite mtETC. Downregulation of these transcripts appears to be a common consequence of genetic ablation of PfMRPs in *P. falciparum*. Several proteins with unassigned functions also had reduced transcript levels following late phase KD. Analysis of these proteins and their tentative functional assignments are discussed below (Table 1). A heatmap of the additional 118 downregulated transcripts along with their GO term designations are shown in Figure S7.

**Table 1.** Potential novel PfMRPs.

|  | <u>Plasmodium</u><br><u>gene ID</u> | <u>Structure prediction</u> | <u>SMLT ID</u> | <u>Orthologs in</u><br><u>Toxoplasma</u> | <u>Orthologs</u><br><u>in Crypto</u> |
| --- | --- | --- | --- | --- | --- |
| 1 | PF3D7_0619600 | Tetrahymena thermophila mS78 | 6z1p.80.A | no | no |
| 2 | PF3D7_1436700 | Tetrahymena thermophila mS89 | 6z1p.91.A | TGME49_227990 | no |
| 3 | PF3D7_0720600 | Tetrahymena thermophila mL64 | 6z1p.39.A | TGME49_225810 | no |
| 4 | PF3D7_1036600 | Tetrahymena thermophila mL40 | 6z1p.43.A | TGME49_266260 | no |
| 5 | PF3D7_1409700 | Tetrahymena thermophila mL104 | 6z1p.43.A | TGME49_300030 | no |
| 6 | PF3D7_0108100 | Tetrahymena thermophila mL104 | 6z1p.43.A | TGME49_249690 | no |
| 7 | PF3D7_0317900 | Tetrahymena thermophila mL104 | 6z1p.43.A | TGME49_229620 | no |
| 8 | PF3D7_1004900 | Tetrahymena thermophila mL105 | 6z1p.44.A | TGME49_254230 | no |
| 9 | PF3D7_0728300 | Trypanosoma brucei mS23 | 6hiv.45.A | TGME49_227860 | no |
| 10 | PF3D7_1309800 | Trypanosoma brucei mS34 | 6hiv.53.A | TGME49_288990 | no |
| 11 | PF3D7_1214000 | At3g22300/P42797 (Plant mitochondrial<br>ribosomal protein uS10) | 6xyw.46.A | TGME49_235730 | no |
| 12 | PF3D7_0111200 | At1g07830/F24B9_7 (Plant mitochondrial<br>ribosome uL29m) | 6xyw.25.A | TGME49_266840 | no |
| 13 | PF3D7_0313200 | 28S Human mitochondrial ribosomal<br>protein S22 | 6nu2.69.A | TGME49_309390 | no |

The parasite’s response to the KD of each MRP also included upregulation of other transcripts. 177 upregulated transcripts overlapped between the late phase responses to the PfRSM22 KD and the PfMRPL23 KD (Figure 5C). Interestingly, only 2 of the 177 upregulated transcripts were related to the parasite mitochondrion; a putative ABC transporter (PF3D7_1352100; ATM-1 like) and a DNL-type Zinc finger protein (Pf3D7_1420300; HSP70-3 co-chaperone). Both genes are non-mutable as assessed by the large-scale mutagenesis study carried out using the PiggyBac transposon (45). A possibility exists that increased expression of these two proteins could be part of a feedback response to mitochondrial dysregulation. Apicoplast chaperone protein DnaJ (Pf3D7_0409400) was also slightly upregulated on day 6 following PfRSM22 KD and day 4 after PfMRPL23 KD (Figure S6B). The upregulation of some of these transcripts as the parasite approaches its demise may indicate attempts to maintain a certain level of homeostasis. Figure S8 provides a heatmap of 175 upregulated transcripts and related GO term pathways affected outside the parasite mitochondrion.

### Late effects of PfRSM22 and PfMRPL23 KD on transcripts of mitoribosomal components

The transcriptomic profiles of parasites at day 6 and day 4 in the PfRSM22 and PfMRPL23 KD lines, respectively, were different from those seen at earlier times following KD. 28 out of the 43 annotated PfMRPs were downregulated on day 4 following PfMRPL23 KD, with the exception of 3 SSU PfMRPs (S9, S11 and S12) that were upregulated (Figure 6A, blue). The response of PfRSM22 KD on day 6 included downregulation of 13 PfMRPs (7 SSU and 6 LSU) (Figure 6A, orange). Yeast and mammalian orthologues of 9 out of these 13 PfMRPs belong to the categories of early and intermediate assembly proteins in the process of mitoribosome complex formation (33). Interestingly, only 3 PfAPRs (L9, L18, L21) were downregulated upon PfRSM22 KD and PfMRPL23 KD at later time points (Figure S6B).

**Figure 6.**
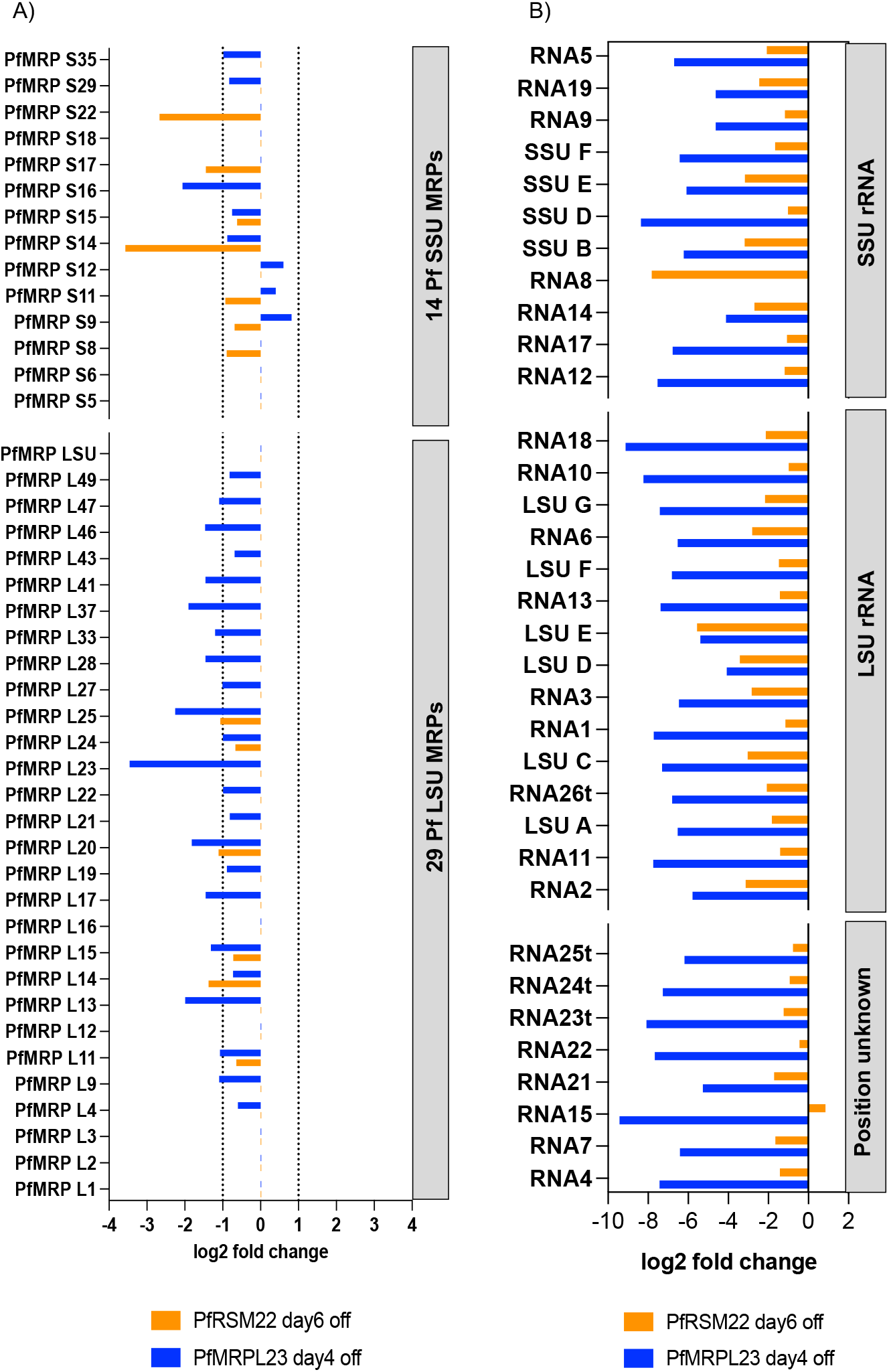
Late effects of PfRSM22 and PfMRPL23 KDs on transcripts of mitoribosomal components.

Among mt rRNAs, 34 out of 39 transcripts were downregulated following the PfRSM22 and PfMRPL23 KDs at the late time points, except for a slight increase in transcription of RNA 15 (Pf3D7_MIT01200) upon PfRSM22 KD (Figure 6B). While the early phase of PfMRPL23 KD on day 2 had slight upregulation of 24 mt rRNA transcripts (Figure 3B), the late phase of PfMRPL23 KD caused a severe downregulation of most mtRNA transcripts. RNA 8 (Pf3D7_MIT04200), an mt SSU rRNA in domain II (Figure S5), was exceptionally downregulated upon PfRSM22 KD but remained undetected in PfMRPL23 KD even at early as well as later time points. As shown in the early phase of KD, 5 mt rRNA transcripts (RNA16, RNA20, SSUA, RNA27t, LSUB) remained undetected in the late phase of KD in both parasite lines.

### Potential novel *Plasmodium falciparum* MRPs

Studying mitoribosomes in *Plasmodium* is hindered by multiple biological and technical challenges of isolating relatively pure mitochondria in tractable quantity. As a result, we have a limited understanding of this divergent ribosome in *Plasmodium spp*. In addition, a large fraction of the *Plasmodium* proteome consists of proteins of unknown function, including 80 proteins included in our provisional mitochondrial proteome (in total of 475). Out of these 80 *P. falciparum* mitochondrial conserved proteins of unknown functions, we found that 71 were differentially regulated upon PfRSM22 and/or PfMRPL23 KD (p < 0.05) giving credence to our bioinformatically derived mitochondrial proteome. Searching for potential roles of these mitochondrial proteins of unknown function, we performed structural modeling using Swiss model (https://swissmodel.expasy.org/). Based on the modeling results, 16 of the differentially regulated mitochondrial proteins of unknown function exhibited structural similarity to known mitoribosomal proteins from other organisms. Transcripts for 13 out of these 16 proteins were significantly regulated following PfRSM22 and/or PfMRPL23 KD (Table 1). Eight out of 13 proteins of unknown function had predicted structural similarity to mitoribosomal proteins from another alveolate, *Tetrahymena thermophilia*(23). Two proteins were structurally similar to MRPs from *Trypanosoma brucei*, two were similar to plant MRPs and one to a human MRP. As expected, the potential PfMRPs listed in Table 1 lack orthologues in *Cryptosporidium parvum*, an apicomplexan genus that has lost mtDNA and, hence, lacks mitochondrial translation. Further studies would need to be performed to determine whether these proteins are divergent or lineage-specific *Plasmodium* MRPs.

### Conclusions

In this study, we have reported the importance of a putative mitoribosome associated protein (PfRSM22) and a ribosomal protein of bacterial origin (PfMRPL23) in the asexual development of *P. falciparum* malaria parasites. We used RNA sequencing as a tool to understand the parasite’s response in regulating gene expression upon KD of PfRSM22 and PfMRPL23. To our knowledge, this is the first report highlighting a potential role of PfRSM22 in mitoribosomal assembly and affecting related downstream pathways leading to parasite death. PfMRPL23 KD on the other hand has a more immediate impact on parasites’ metabolic pathways that is detrimental to the parasite one IDC sooner than PfRSM22 KD. The data also uncovers global and mitochondrial specific transcriptomic changes over 6 days following PfRSM22 KD and 4 days following PfMRPL23 KD. In addition to that, we identified 13 conserved mitochondrial proteins of unknown function that are potential new MRPs in *P. falciparum*.

## EXPERIMENTAL PROCEDURES

### 1. Parasite culture

The D10 strain of *P. falciparum* was cultured in O^+^ human RBCs (Interstate Blood Bank, TN) supplemented with RPMI1640 medium along with 0.3-0.5% Albumax I (Invitrogen), 2g/L sodium bicarbonate, 10 mg/L hypoxanthine (Thermo Fisher Scientific), 15 mM HEPES (Millipore Sigma) and 50 mg/L gentamycin (VWR). Parasite culture was maintained at 37°C with conditions previously described (18).

### 2. Plasmid construction and transfection

The plasmid carrying the TetR-DOZI-aptamer elements for conditional KD (pMG75) was kindly provided by Dr. Jacquin Nile’s group from MIT. Primers used for cloning the homologous regions of interest into the pMG75 vector are listed in Table S1. Guide RNA (gRNAs) were designed using the Eukaryotic Pathogen CRISPR guide RNA Design Tool (http://grna.ctegd.uga.edu). All gRNAs were independently cloned into the MCas9-yDHOD (-) or NF-Cas9-yDHOD (-) vector modified from the pAll-In-One (pAIO) vector provided by Dr. Joshua Beck from Iowa State University (19). The cloning procedure of gRNA was carried out using DNA Assembly Master mix (New England Biolabs, Inc). Wild type D10 *P. falciparum* was transfected at the early ring stage with linearized template vector (pMG75, 50 μg) digested with EcoRV (NEB) overnight along with circular gRNA plasmids (40 μg each). Post transfection, the culture was maintained in media supplemented with 250 nM anhydrotetracycline (aTc, Millipore Sigma) for first two days, followed by blasticidin (BSD) + aTc media. The transfected parasite genotype was confirmed by PCR analysis. Details of cloning procedure is shown in Supporting Information.

### 3. Immunofluorescence Assay (IFA)

A volume of 50 μL parasitized RBCs at trophozoite stage (~5% parasitemia) was collected for IFA. The parasites were labeled with 50 nM Red Mito-Tracker CMXRos (Thermo Fisher Scientific) for 20 mins. This was followed by three washes using 1X PBS before fixing the cells with 4% paraformaldehyde, 0.0075% glutaraldehyde for 1hr at 37 °C on a rotator. The fixed cells were processed as described in our previous publication and visualized under Nikon Ti microscope (19). The anti-HA antibody (Sc-7392, Santa Cruz) and secondary antibody (A11029, Invitrogen) were used in 1:300 dilution.

### 4. Parasite Growth Assay

The endogenously tagged parasite lines were tightly synchronized with two rounds of 0.5 M alanine/10 mM HEPES (pH 7.4) at the ring stage. At the trophozoite stage, synchronized parasites with high parasitemia (8-10%) were washed 3 times with regular RPMI to remove aTc. Washed cells were split 1:10 with new RBCs and were equally transferred to two T25 flasks to allow the parasite growth with and without aTc (250 nM). Parasite morphology and growth rates were monitored in 48 h intervals in Giemsa-stained thin blood smears. Parasitemia was counted per 1000 RBCs under Leica light microscope. Every 48h post KD, the culture was split 1:5 over several IDCs, to collect protein samples for Western blot analysis. Growth curve data was analyzed using GraphPad Prism 9.

### 5. Western blotting

Parasite culture was harvested at the trophozoite stage and treated with 0.1% saponin/PBS supplemented with 1X protease inhibitor (PI) (APEX Bio). The saponin treated parasites were washed with 1X PBS and PI to remove residual hemoglobin. The parasitized pellet was mixed thoroughly with 10 volumes of 3% SDS/60 mM Tris-HCl (pH 7.5) and solubilized overnight at 4°C on a rotator. The sample was spun down at 12,000 rpm for 10 mins to acquire supernatant for SDS-PAGE. Protein concentration was determined by the Pierce BCA Protein Assay (23227, ThermoFisher). Post electrophoresis, protein was transferred to methanol activated PVDF membrane (0.45 μm, HybondTM) at 23V overnight at 4°C. The membrane was blocked with 5% milk/PBS for 2 h before incubation with mouse monoclonal anti-HA antibody (Sc-7392, Santa Cruz) at 1:10,000 for overnight at 4°C. Horseradish peroxidase (HRP) conjugated secondary goat anti-mouse antibody (Cat: 62-6520, Thermo Fisher Scientific) at 1:10,000 was used for 4 h at room temperature. The membrane was incubated with Pierce ECL Western Blotting Substrate and visualized using the BioRad imager. The membrane was further probed with 1:10,000 rabbit anti-PfExp2 primary antibody followed by 1:10,000 HRP conjugated secondary anti-rabbit antibody (Thermo Fisher) for 1h, before it was developed to show loading controls.

### 6. RNA extraction and library construction

D10-PfRSM22-3HA and D10-PfMRPL23-3HA parasite lines were tightly synchronized and grown in T175 flasks in presence of aTc. At the trophozoite stage (day 0), the parasites were washed and split into new RBCs to start KD. The parasites were harvested at late trophozoite stage on day 2, day 4 and day 6 by saponin lysis (0.05% in PBS). Day 2 aTc plus parasite culture was harvested which served as controls. The saponin lysed parasite pellets were immediately dissolved in 6 volumes of TriZol (Invitrogen) and kept frozen at −80°C until to be thawed on ice for RNA isolation. For each sample, 1/5^th^ volume of chilled chloroform was added, followed by a spin in 4°C at 4000 rpm for 10 mins. The top aqueous layer was transferred to a fresh chilled tube without disturbing the middle buffy coat. 100% ethanol was added to the aqueous layer in a 1:1 ratio. This mixture was subjected to RNA isolation using a kit according to the manufacture’s protocols (Zymo research). The concentration of eluted RNA was determined using Nanodrop (Thermo Fisher). Quality of eluted RNA was tested by running 1μg of RNA on agarose gel at 4°C to visualize intact rRNA. Intact rRNA bands were confirmed indicating no obvious RNA degradation.

Isolated RNA samples were sent to Novogene Co for sequencing where the samples were further quantified using a qubit fluorometer (Thermo Fisher, Waltham, MA) and the quality was scored using a bioanalyzer (Agilent Technologies, Santa Clara, CA). Samples with RIN 8.0 and above were selected for library preparation. Briefly, 1μg of input RNA was used for RNA library preparation using the NEB Next Ultra II RNA Library Prep kit for Illumina (E7775L; NEB). Prepared libraries were quantified using real time PCR and bioanalyzer. Libraries of 3 nM and above concentration were loaded on to NovaSeq 6000 S4 Reagent Kit using a paired-end 150 kit (20012866; Illumina, San Diego, CA). Each sample was sequenced to a depth of at least 19.5 million reads. The number of clean reads obtained from each sample is listed in Table S5.

### 7. RNA sequencing and data analysis

The fastq files were generated using HWI-ST1276 instrument. Quality of reads was checked using FastQC tool and later processed to remove the adapter sequences and to perform base quality control check (46). Quality metrics and error rate were transformed using Phred score with highest stringencies (47). Fastp processed files were aligned to the most updated *P. falciparum* reference genome using HISAT2 software (v2.0.5) (48). Exons comprised of more than 90% of the sequences. RPKM counts of each gene was counted using Stringtie (v1.3.3) (49). Differential expression between control and KD samples was calculated using DEseq2 R package (v1.20.0) (50). Samples were processed as duplicates and analyzed individually for differential expression. The adjusted p value was calculated after correction of p value using Benjamini and Hochberg method. A p-value < 0.05 considered to obtain significant DESeq2 values. Gene ontology (GO) and KEGG (Kyoto Encyclopedia of Genes and Genomes, http://www.kegg.jp/) enrichment analysis of differentially expressed genes was performed using clusterProfiler R package (v3.8.1) (51). GSEA analysis was performed using gsea v3.0. PlasmoDB search tools were used to make lists of gene IDs mentioned in Table S4. Volcano plots, heat maps and other graphs were made on GraphPad Prism 9.

## Data availability statement

The FASTq sequence files of PfRSM22 and PfMRPL23 from RNA sequencing reported in this paper are deposited in NCBI under the project number PRJNA801275. All additional data are contained within this manuscript and in the Supporting Information.

## Acknowledgements

We are grateful for the constant support from members of the Center for Molecular Parasitology, Department of Microbiology and Immunology at Drexel University College of Medicine. We are thankful to Dr. Aishwarya Iyer (University of Alberta, Canada), Dr. Frank Bearof and Dr. Joshua Mell (Drexel University College of Medicine) for their guidance in RNA sequencing data analysis. We also thank Dr. Jacquin Niles (MIT) for the TetR-DOZI-aptamer plasmid, Dr. James Burns (Drexel University College of Medicine) for Exp2 antibody and Dr. Joshua Beck (Iowa State University) for the CRISPR/Cas9 plasmid. We thank vEuPathDB (https://veupathdb.org) for providing resources for data analysis. The work was supported by NIH/NIAID grants to Dr. Hangjun Ke (K22AI127702) and Dr. Akhil Vaidya (AI028398).

## Author contributions

HK and ABV designed and guided the experiments. SW and LL designed plasmid construction and generated parasite lines. SW performed experiments including RNA isolation and analyzing sequencing results generated by Novogene Co Inc. MWM and SW conducted bioinformatic analysis. SW wrote the manuscript, which was edited by MWM, ABV and HK.

## FOOTNOTES

* This work was supported by an NIH/NIAID career transition award (K22AI127702) to Dr. Hangjun Ke and an NIH/NIAID R01 grant to Akhil B. Vaidya (AI028398).

The abbreviations used are:

PfRSM22: Plasmodium falciparum mitochondrial ribosomal associated protein of small subunit
METTL17: methyltransferase like protein 17
SAM: S-adenosyl-L-methionine dependent methyltransferase
mitoribosomes: mitochondrial ribosome
MRP: mitoribosomal protein
mtETC: mitochondrial electron transport chain
mtDNA: mitochondrial DNA
cyt: cytochrome
cox1: cytochrome oxidase subunit I
cox3: cytochrome oxidase subunit III
TetR: tetracycline repressor
DOZI: development of zygote inhibited
cryo-EM: cryo-electron microscopy
gRNA: guider RNA
aTc: anhydrotetracycline
IDC: intraerythrocytic cycles
KEGG: Kyoto Encyclopedia of Genes and Genomes
GO: Gene Ontology
RPKM: Reads Per Kilobase Million
GSEA: Gene Set Enrichment Analysis

