## Supplementary Information for "Transcriptional changes in *Plasmodium falciparum* upon conditional knock down of mitochondrial ribosomal proteins RSM22 and L23"

1. Plasmid construction for knocking down PfRSM22 and PfMRPL23.
2. Figure S1. PfRSM22 protein domain and amino acid sequence alignment.
3. Figure S2. PfMRPL23 protein domain and amino acid sequence alignment.
4. Figure S3. Endogenous tagging of PfRSM22_3HA and PfMRPL23_3HA using CRISPR/Cas9.
5. Figure S4. Downregulation of PfRSM22 and PfMRPL23 mRNA post aTc removal.
6. Figure S5. Secondary structure of Pf mt rRNA fragments and their likely positions in the modeled SSU and LSU.
7. Figure S6. Early and late effects of PfRSM22 and PfMRPL23 KD on apicoplast related transcripts.
8. Figure S7. Downregulated non-mitochondrial transcripts common between PfRSM22 and PfMRPL23 KDs in the late phase.
9. Figure S8. Upregulated non-mitochondrial transcripts common between PfRSM22 and PfMRPL23 KDs in the late phase.
10. Table S1. List of primers and oligoes used in this study.
11. Table S2. Read count files of the PfRSM22_3HA line (aTc ON and day 2, day4, day 6 aTc OFF) and the PfMRPL23_3HA (aTc ON and day 2, day 4 aTc OFF).
12. Table S3. KEGG and Gene ontology (GO) term list of significantly regulated genes upon KD of PfRSM22 and PfMRPL23.
13. Table S4. List of genes in Venn diagrams shown in Figure 5A and B.
14. Table S5. RNA sequencing read depth.

### 1. Plasmid construction for knocking down PfRSM22 and PfMRPL23

All primers and oligos were purchased from Eurofins Genomics and Genewiz LLC (Table S1). DNA fragments were amplified using high fidelity DNA polymerases (New England Biolabs®, Inc) and confirmed by sequencing (Genewiz LLC). Transformation of pMG75 related plasmids was performed using Stable competent *E. coli* cells (New England Biolabs®, Inc) and bacteria were grown at 30°C to maintain the stability of 8 aptamer repeats. Transformation of Cas9 related plasmids was performed using DH5-alpha electrocompetent *E. coli* cells and bacteria were grown at 37°C.

The pMG75 vector [1] was used to modify the genomic locus of PfRSM22 (Pf3D7_1027200). Briefly, the original pMG75noP-ATP4-8apt-3HA plasmid was linearized by *Afl*II and *BstE*II*,* to remove the ATP4 inserts. PfRSM22 5'HR region was amplified from wild type *P. falciparum* (WT) genomic DNA using primers (P2+P3) giving a PCR product of 868 bp. The 3'HR region was downstream of the stop codon and was amplified using primers (P4 + P5) giving a PCR product of 601 bp in length. The three pieces including the linearized vector, PfRSM22 5'HR and PfRSM22 3'HR were joined together using NEB HiFi DNA assembly master mix. The gRNAs of PfRSM22 were present at the end of 5'HR where Cas9 would introduce cut to create the expected modification in the transgenic parasites. To avoid repetitive cutting, synonymous mutations were introduced in the reverse primer of the 5'HR (P3) within the gRNA region. The modified pMG75 vector was sequenced using primers P11 and P12 to verify 3'HR and 5'HR respectively.

The PfMRPL23 5'HR and 3'HR were cloned to the pMG75 vector one after the other. Briefly, the pMG75 vector containing mt-DNA polymerase I (Pf3D7_0625300) was digested with BstEII and SacII to remove mt-DNA pol 3' HR. PfMRPL23 3' HR was amplified from WT parasite DNA using 3'HR primers flanked by BstEII at its reverse primer and SacII at its forward primer (P16 +P17). The amplified 3'HR (1000 bp) was digested with SacII and BstEII and inserted into the digested pMG75 vector. Presence of PfMRPL23 3'HR and absence of mt-DNA pol 3'HR was confirmed by PCR amplification followed by sequencing analysis. The pMG75 vector carrying PfMRPL23 3'HR was digested with SacII and SalI to remove mtDNA pol 5'HR. PfMRPL23 5' HR was amplified using primers P14+P15. The amplified product was 853 bp in length and was digested with SacII and SalI and inserted into the pMG75 vector carrying PfMRPL23 3'HR, resulting in the final plasmid pMG75-PfMRPL23-3HA-8apt. The modified pMG75 vector was sequenced using primers P11 and P12 to verify 3'HR and 5'HR respectively.

gRNAs were designed using Eukaryotic Pathogen CRISPR guide RNA Design Tool (http://grna.ctegd.uga.edu/). High efficiency score gRNAs having no off-target matches were selected and PfRSM22 gRNAs (P7 and P9) were cloned in No flag (NF)-Cas9-yDHOD (-) construct. Briefly, NF-Cas9-yDHOD (-) vector [1] was digested with EcoRI and joined with oligos P7 and P9 individually by NEB HiFiDNA assembly. Correct infusion of PfRSM22 gRNA1-NF-Cas9-yDHOD (-) was confirmed by PCR using primers P8+P23 whereas PfRSM22 gRNA2 infusion was confirmed by PCR using P10+P23. The inserted gRNAs were further confirmed by sequencing analysis. PfMRPL23 gRNAs were cloned into the M-Cas9-yDHOD (-) vector that still had a flag tag present in Cas9 [2]. The M-Cas9-yDHOD(-) vector digested with EcoRI was joined with oligos P19 and P21 individually by NEB HiFiDNA assembly. Fusion of gRNAs into M-Cas9-yDHOD was verified by PCR using primers P20+P23 (gRNA1) or P22+P23 (gRNA2) and sequencing.

2. **Figure S1. PfRSM22 protein domain and amino acid sequence alignment.** (A) The green bars represent the length of each protein, while the brown and pink bars indicate the InterPro RSM22 family and methyltransferase domain regions within each protein, respectively (drawn to scale). The numbers represent the length of each protein in amino acids. Protein families and domains were identified via InterPro search: (https://www.ebi.ac.uk/interpro/search/sequence). RSM22-like proteins are shown with Uniprot identifier: *Bradyrhizobium japonicum* Q89SB0, *Plasmodium falciparum* Q8IJD1, *Toxoplasma gondii* S8GCN9, *Tetrahymena thermophila* Q22CT3, *Homo sapiens* P82650, *Caenorhabditis elegans* P91862, *Saccharomyces cerevisiae* P36056, *Arabidopsis thaliana* Q8GW63, *Trypanosoma brucei* Q385R2. (B) Protein sequence alignment of RSM22 orthologues from the organisms listed above. Intensity bars indicating the quality represent conservation of amino acids across listed organisms.

**(A)**

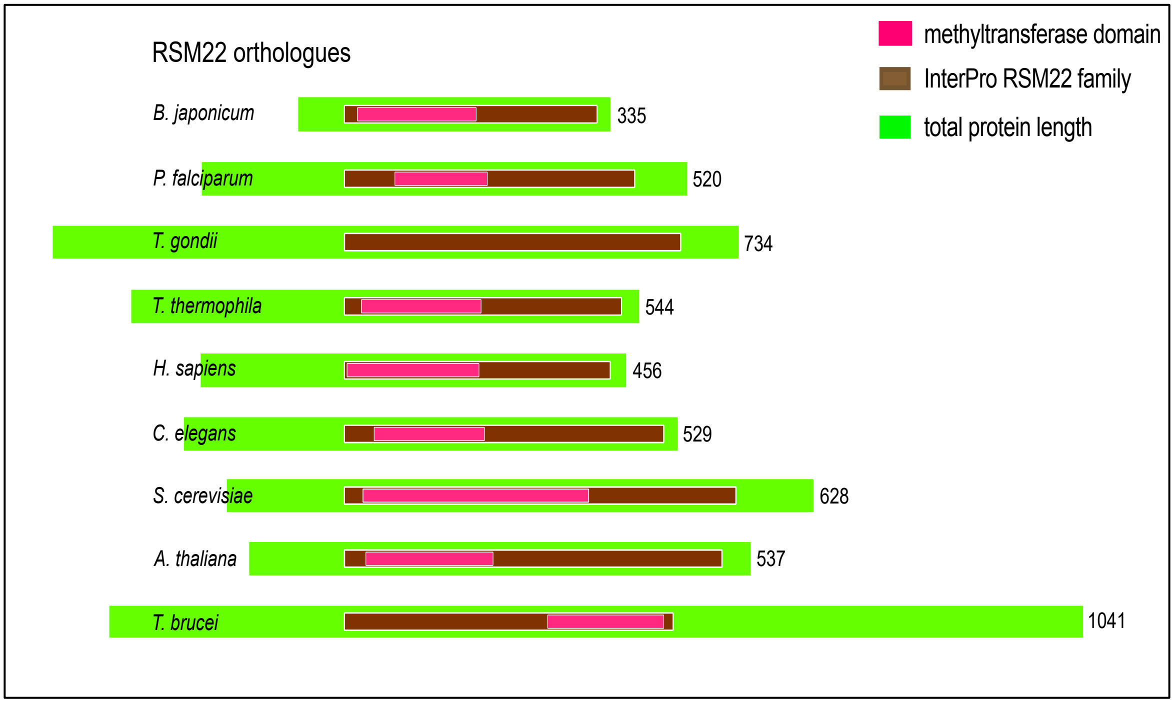

(B)

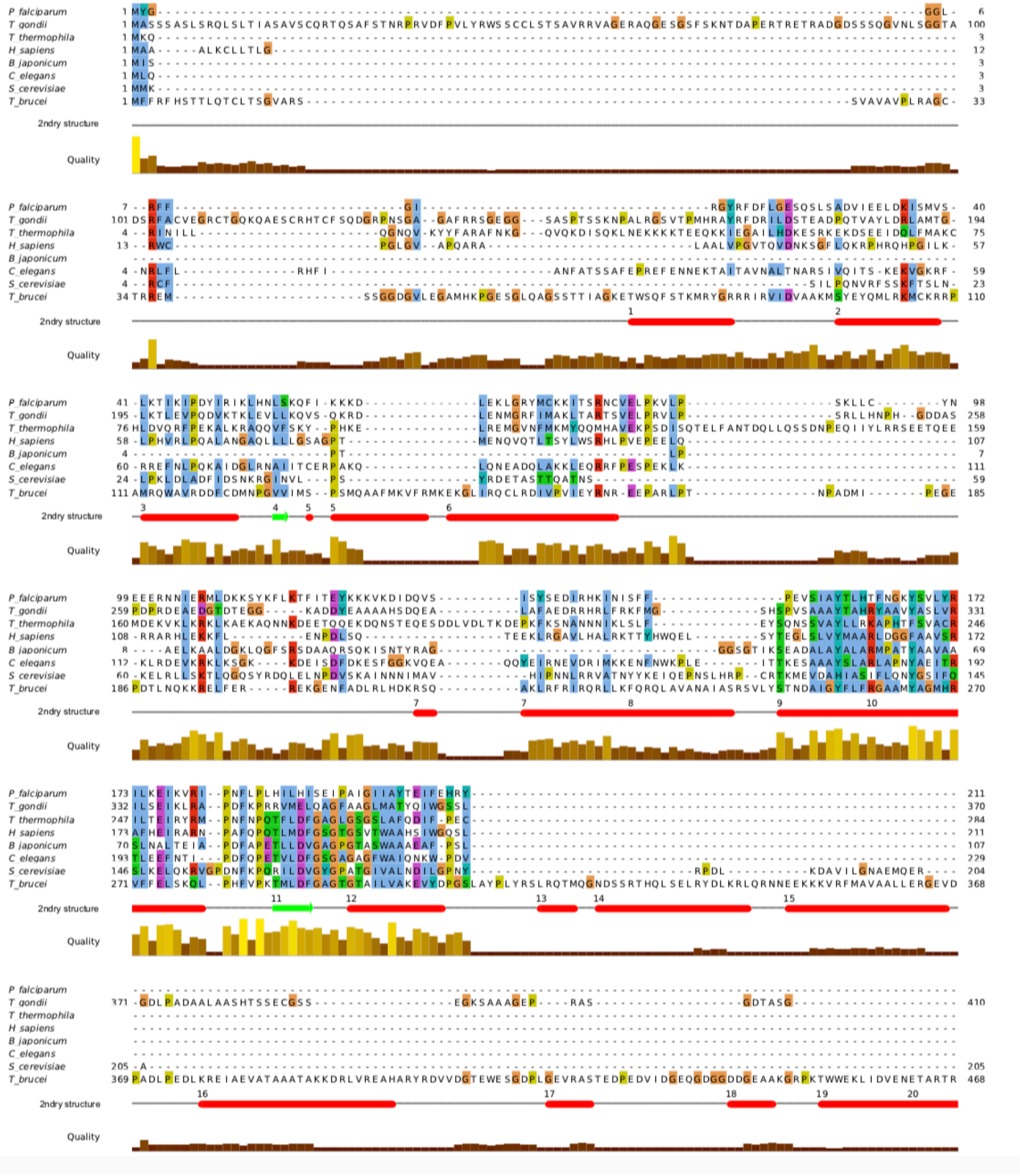
P falciparum
T gondii
T thermophila
H sapiens
B japonicum
C elegans
S cerevisiae
T brucei
2ndry structure
Quality

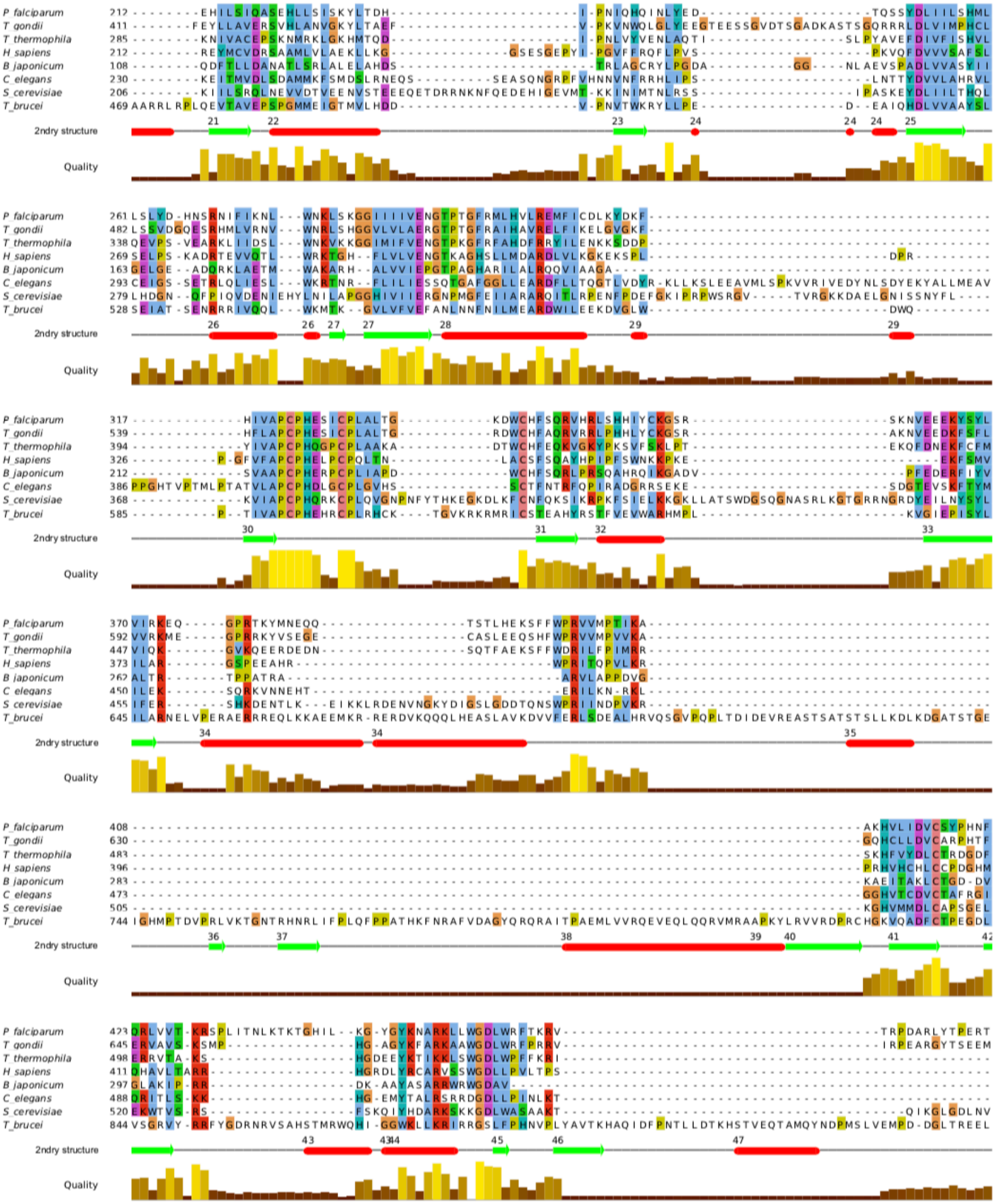
P falciparum
T gondii
T thermophila
H sapiens
B japonicum
C elegans
S cerevisiae
T brucei
2ndry structure
Quality

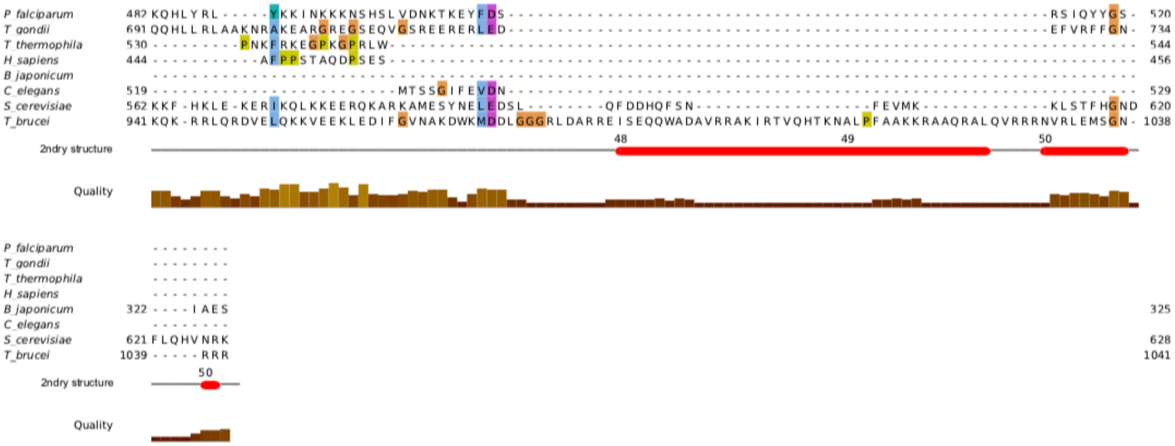
P falciparum
T gondii
T thermophila
H sapiens
B japonicum
C elegans
S cerevisiae
T brucei
2ndry structure
Quality

3. **Figure S2. PfMRPL23 protein domain and amino acid sequence alignment.** (A) Green and brown bars represent the length of each protein and uL23 protein family region within each protein, respectively. The numbers represent the length of each protein in amino acids. Protein families were identified via InterPro search(https://www.ebi.ac.uk/interpro/search/sequence). uL23 proteins are shown with Uniprot identifier: *Escherichia coli* P0ADZ0, *Rickettsia prowazekii* Q9ZCQ7. *Saccharomyces cerevisiae* P32387, *Homo sapiens* Q16540, *Plasmodium falciparum* Q8I532, *Toxoplasma gondii* A0A125YSK9, *Trypanosoma brucei* Q387G3, *Arabidopsis thaliana* Q9SMR5, *Tetrahymena thermophila* Q22EY1. (B) Protein sequence alignment of uL23 proteins from organisms listed above. Intensity bars indicating the quality represents conservation of amino acids across listed organisms.

**(A)**

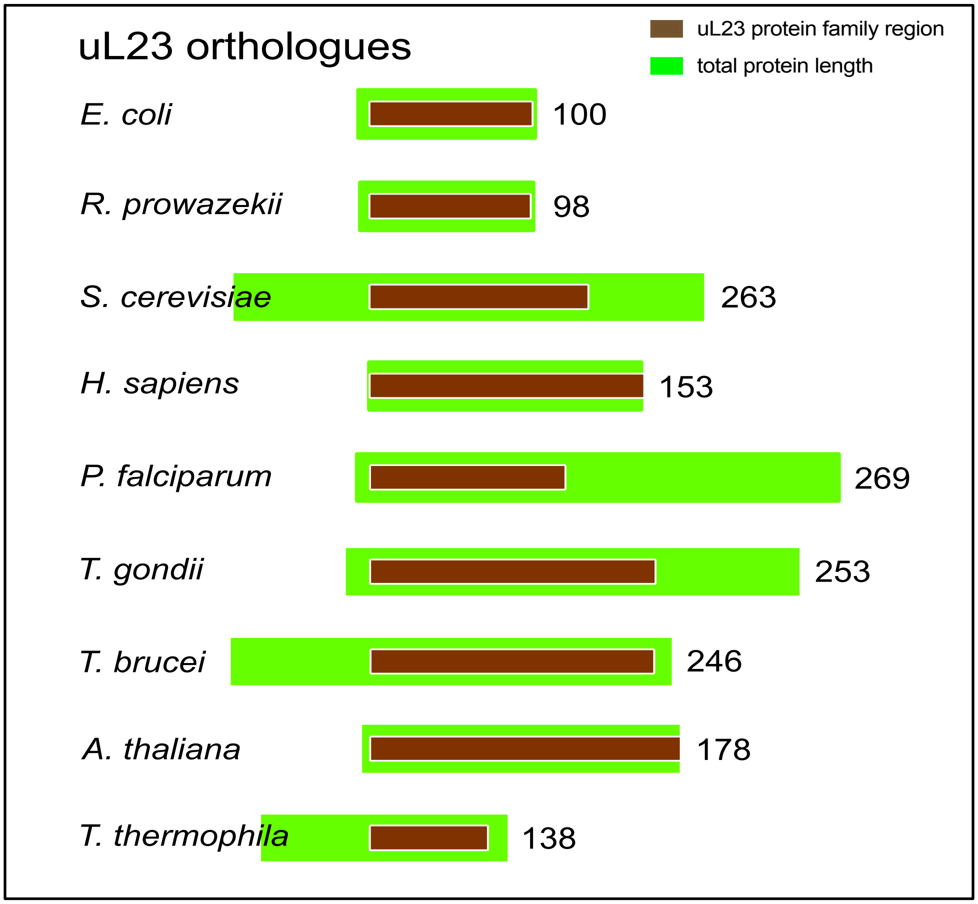

**(B)**

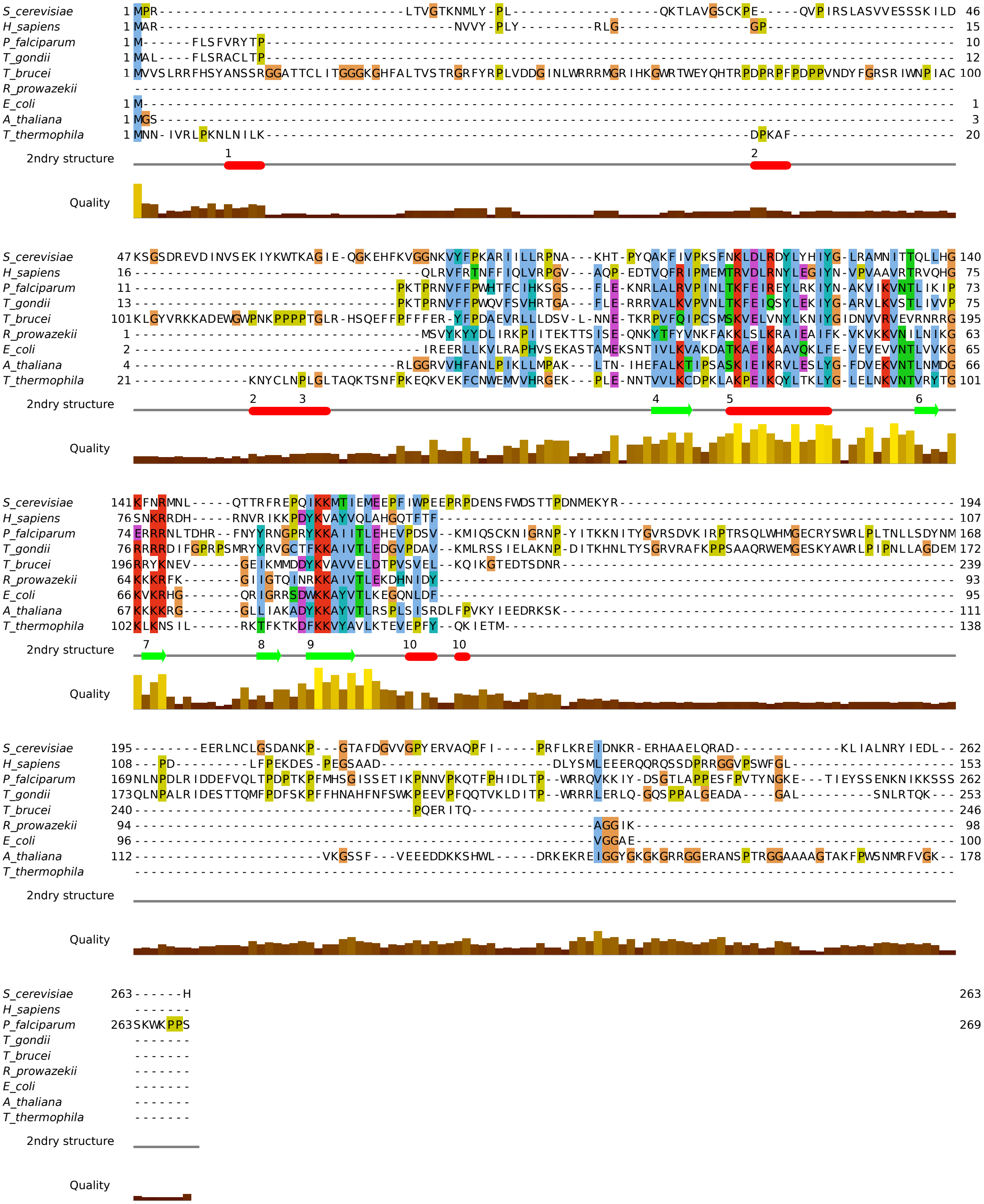
S_cerevisiae
H_sapiens
P_falciparum
T_gondii
T_brucei
R_prowazekii
E_coli
A_thaliana
T_thermophila
2ndry structure
Quality

**4. Figure S3. Endogenous tagging of PfRSM22_3HA and PfMRPL23_3HA using CRISPR/Cas9.** (A) A schematic of endogenous gene modification of PfRSM22 and PfMRPL23 using CRISPR/Cas9 mediated double crossover recombination. The pMG75-TetR-DOZI-8aptamer plasmid was linearized with EcoRV and transfected into WT D10 parasites together with corresponding circular gRNA plasmids. Via double crossover recombination, the gene locus of PfRSM22 or PfMRPL23 was added with a 3xHA tag and 8 aptamer repeats. Position of primers used to verify the parasite genotype in B is mentioned. (B) Genotyping of D10-PfRSM22_3HA and D10-PfMRPL23_3HA parasite lines. DNA gel imaging of PCR products amplified from integrated or WT DNA, representing correct integration of 5' HR and 3' UTR at the expected gene loci. WT gene locus is intact in WT parasites.

A.

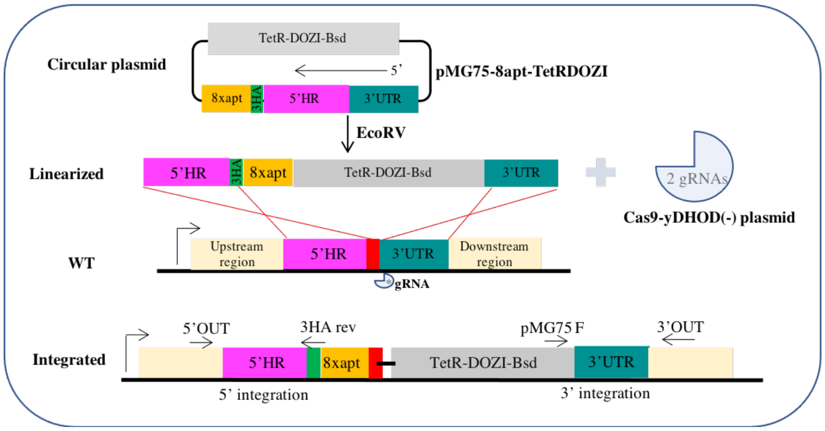


B.

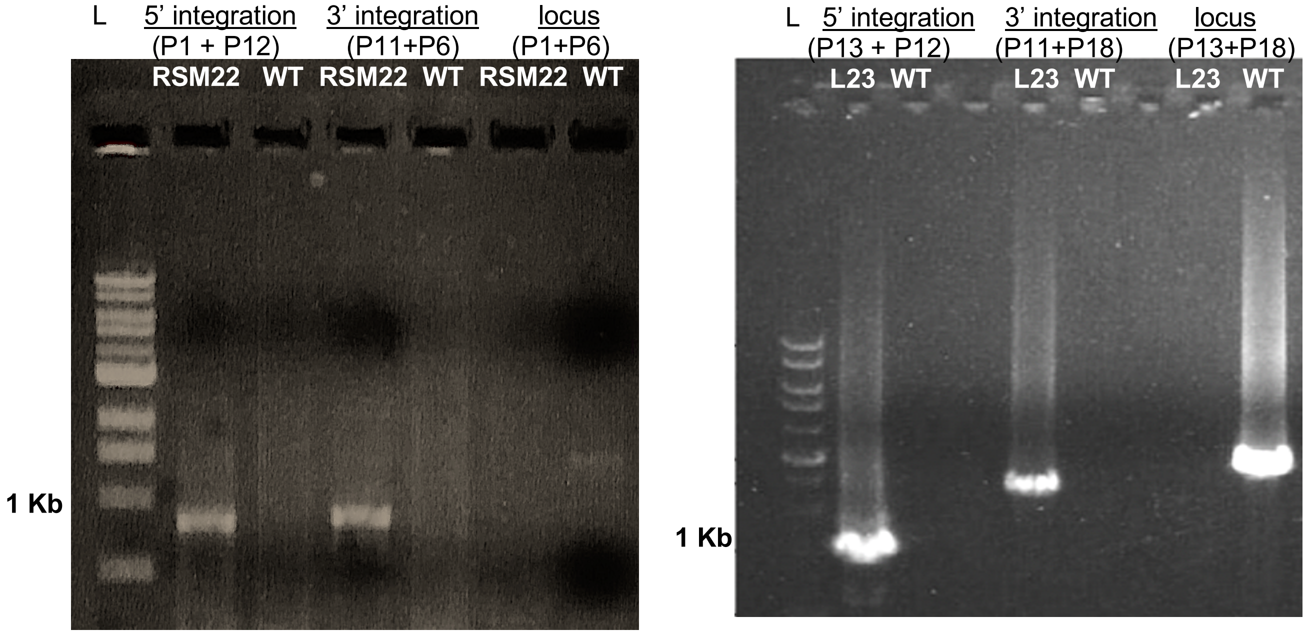

5. **Figure S4. Downregulation of PfRSM22 and PfMRPL23 mRNA post aTc removal.** Total RNA sequencing result showing reduction of PfRSM22 and PfMRPL23 shown in log2fold change compared to PfRSM22 and PfMRPL23 mRNA in presence of aTc. Data shown is the mean of duplicates with $p < 0.05$.

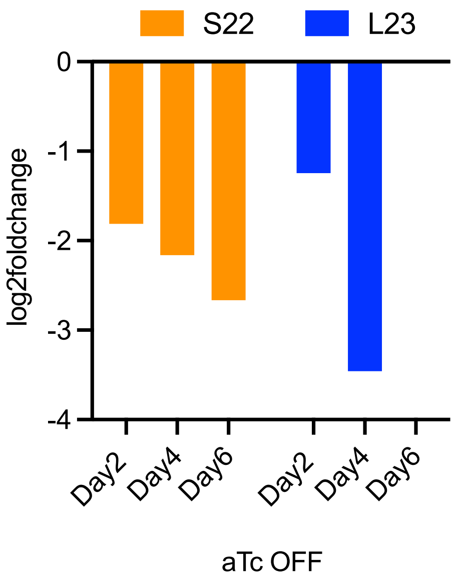

6. **Figure S5: Secondary structure of Pf mt rRNA fragments and their likely positions in the modeled SSU and LSU.** Secondary structures of SSU and LSU Pf mt rRNA fragments represented in light to dark grey color, subdivided into domain I, II, III, IV, V and VI. Transcripts uniquely downregulated upon PfRSM22 KD are represented in orange. No transcript was uniquely regulated upon PfMRPL23 KD. Transcripts differentially regulated upon KD of both PfRSM22 and PfMRPL23 are represented in teal. Purple color represents transcripts that were not detected in this study. The most updated *Plasmodium* mt rRNA map does not include position of 11 mt rRNA transcripts [3].

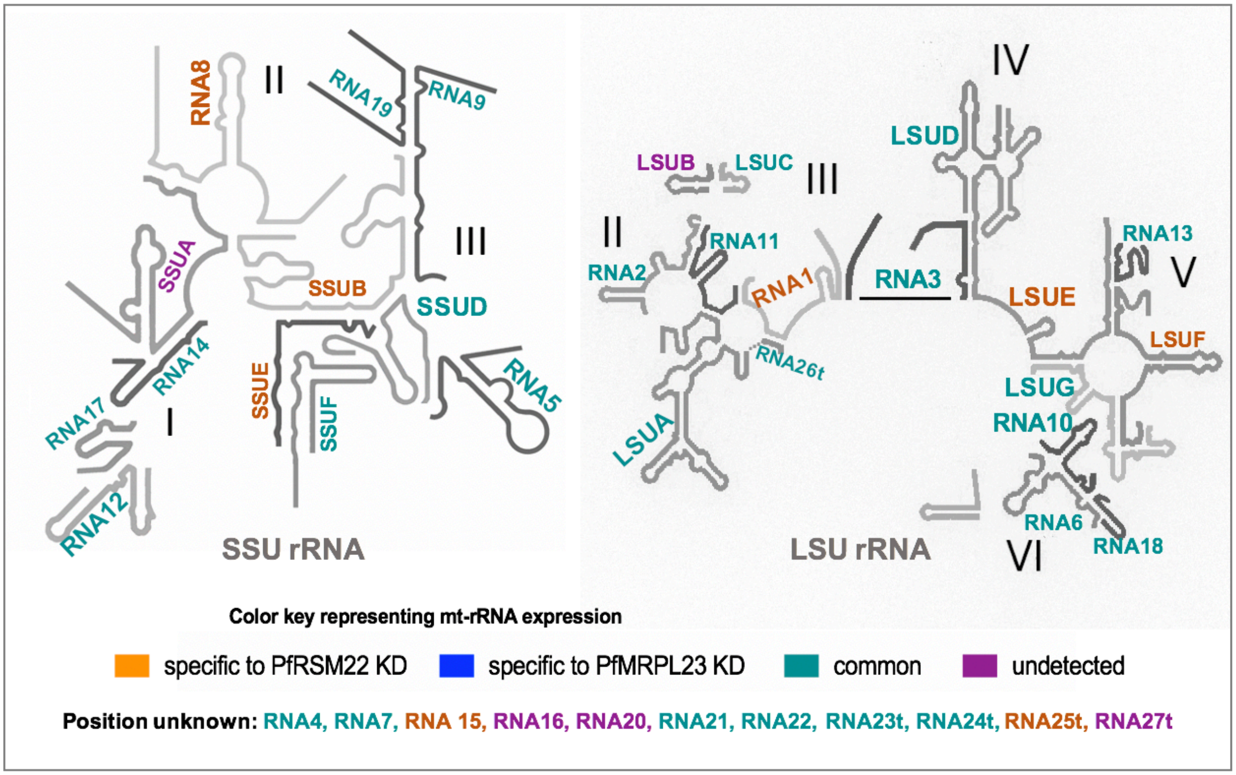

7. **Figure S6: Early and late effects of PfRSM22 and PfMRPL23 KD on apicoplast related transcripts.** (A) Heat map of differentially regulated transcripts common in the early phase of PfRSM22 and PfMRPL23 KD that are likely localized to the apicoplast. (B) Heat map of differentially regulated transcripts common in the late phase of PfRSM22 and PfMRPL23 KD that are likely localized to the apicoplast. The apicoplast proteome was determined previously [4].

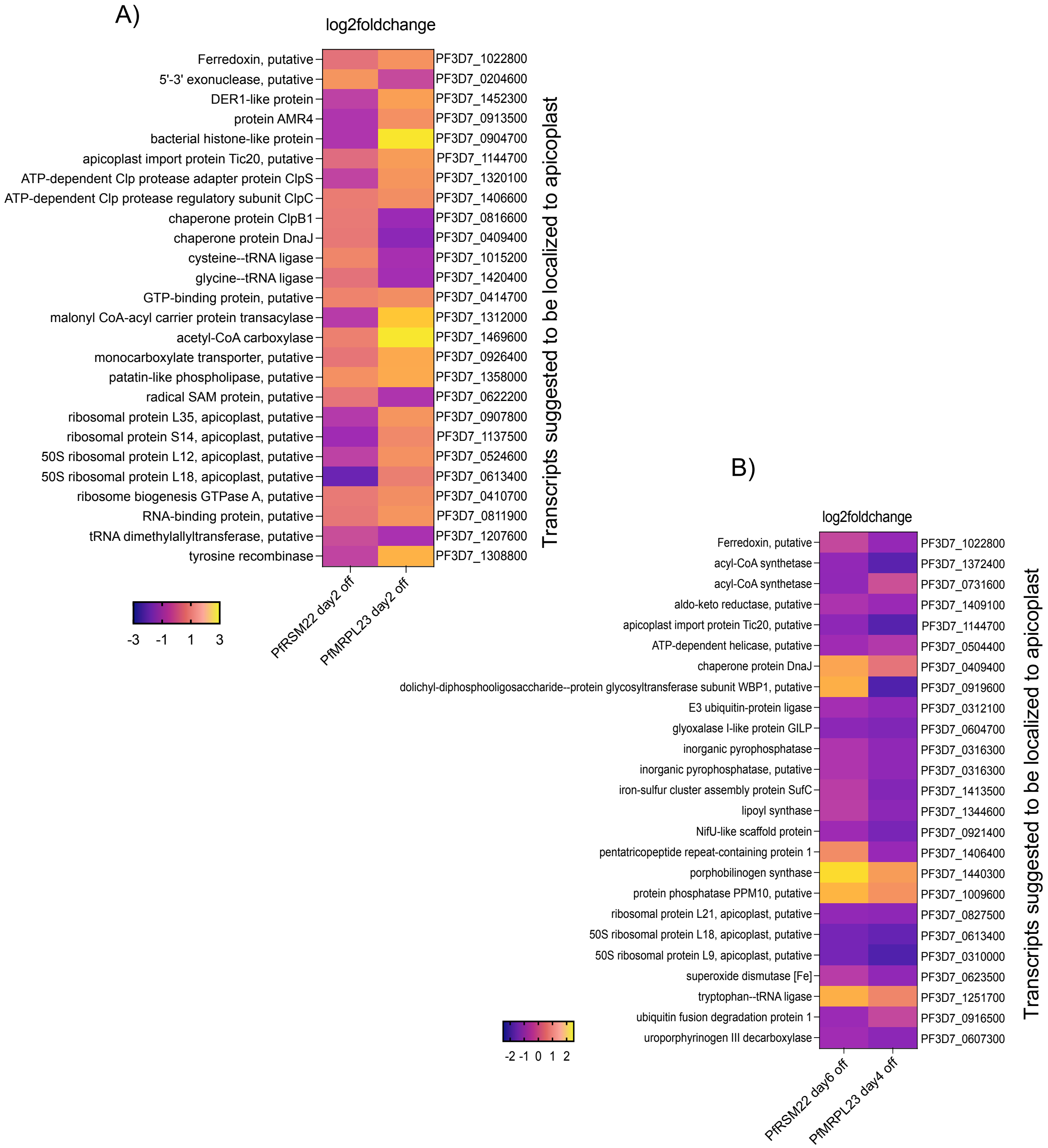

A.

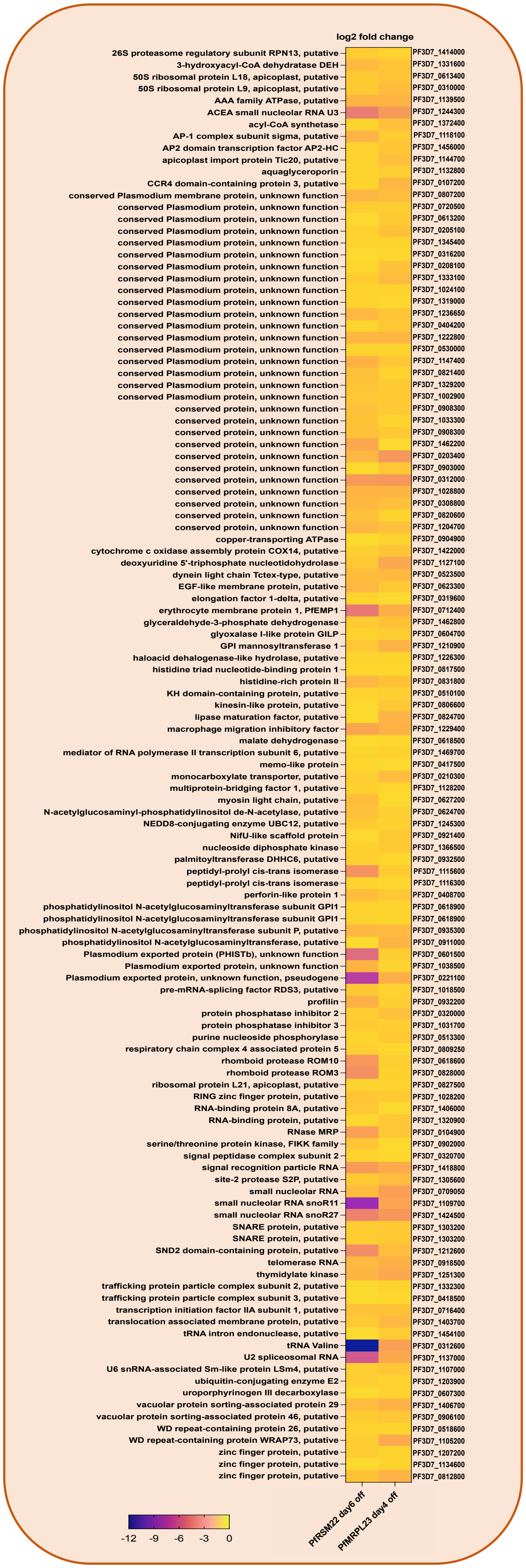


**8. Figure S7: Downregulated non-mitochondrial transcripts common between PfRSM22 and PfMRPL23 KDs in the late phase.** (A) Heat map of downregulated non mitochondrial 118 transcripts in common between PfRSM22 day6off and PfMRPL23 day4off. (B) List of GO term pathways of 118 transcripts generated on g:Profiler web server (https://biit.cs.ut.ee/gprofiler/gost)

B.

| Term name | Term ID |
|---|---|
| membrane lipid biosynthetic process | GO:0046467 |
| membrane lipid metabolic process | GO:0006643 |
| glycolipid metabolic process | GO:0006664 |
| glycolipid biosynthetic process | GO:0009247 |
| liposaccharide metabolic process | GO:1903509 |
| pyrimidine nucleoside triphosphate metabolic process | GO:0009147 |

A.

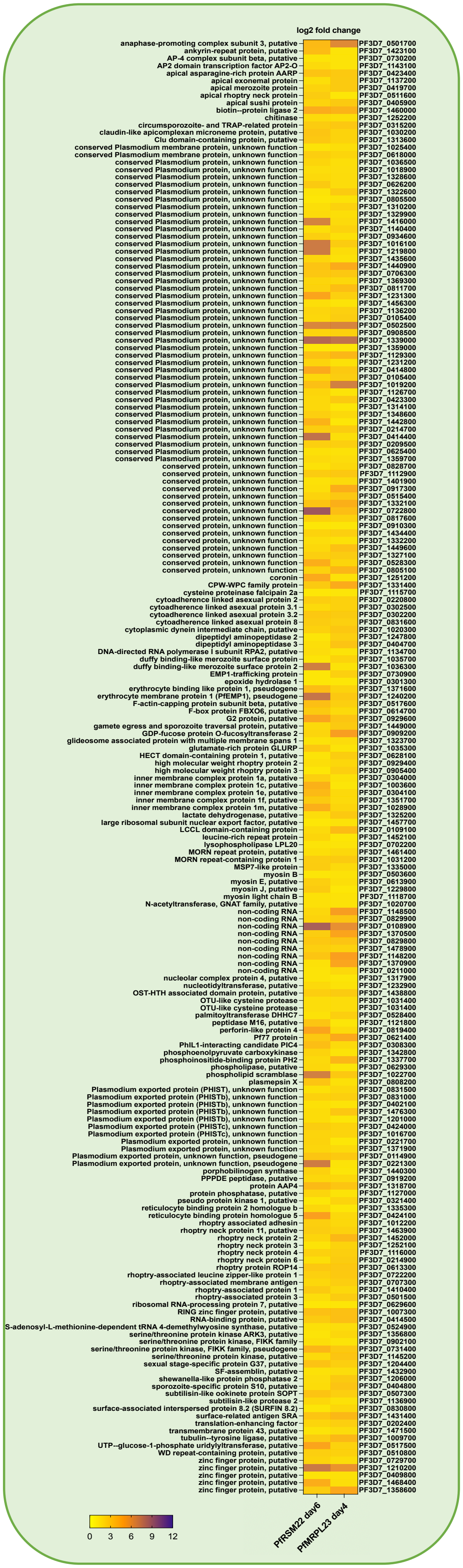


**9. Figure S8: Upregulated non-mitochondrial transcripts common between PfRSM22 and PfMRPL23 KDs in the late phase.** (A) Heat map of upregulated non mitochondrial 175 transcripts in common between PfRSM22 day6off and PfMRPL23 day4off. (B) List of GO term pathways of 175 transcripts generated on g:Profiler web server (https://biit.cs.ut.ee/gprofiler/gost)

B.

B.

| Term name | Term ID |
|---|---|
| entry into host | GO:0044409 |
| movement in host environment | GO:0052126 |
| cytoskeleton-dependent intracellular transport | GO:0030705 |
| cytoskeleton organization | GO:0007010 |
| biological adhesion | GO:0022610 |
| actin filament organization | GO:0007015 |
| actin filament-based process | GO:0030029 |
| actin cytoskeleton organization | GO:0030036 |
| supramolecular fiber organization | GO:0097435 |

**4. Table S1: List of primers and oligoes used in this study.**

| | Primers | Sequence |
|---|---|---|
| P1 | PFRSM22-5FOUT | AGATCACATACCAAATATAC |
| P2 | PFRSM22-5HRFWD | GGCCGCGGGATATCTCCGGAGTAGAGAATGGTACACCCACAG |
| P3 | PFRSM22-5HRREV | AAAATGTTTATCAAACCGGGGGTAACCTGTGATCCATAGTATTGAA TTGATCTAGAATCGAAATATTCCTTAGTCTTGTT |
| P4 | PFRSM22-3UTRFWD | ATGGCCCCTTTCCGGGCGCGCCTTAAGGAATAACACAAGAG GTTATAGTTTA |
| P5 | PFRSM22-3UTRREV | TCCGGAGATATCCCGCGGCCTATTTGATGAGTGCATTATCC |
| P6 | PFRSM22-3FOUT | TAAAATAAGTGTGTGGTGCT |
| P7 | PFRSM22_gRNA1 | CATATTAAGTATATAATATTGTCACGTTCCATACAGTATTAGT TTCAGAGCTATGCTGGA |
| P8 | PFRSM22_gRNA1_N21 | GTCACGTTCCATACAGTATTA |
| P9 | PFRSM22_gRNA2 | CATATTAAGTATATAATATTATATGTGTAGCATATTTCTTGTT TCAGAGCTATGCTGGAA |
| P10 | PFRSM22_gRNA2_N20 | ATATGTGTAGCATATTTCTT |
| P11 | PMG75seqF | CTTTAAATTCATGCAAAAATTTAC |
| P12 | BBHA REV | TGGGCCCGAATTCTCATCATTGTGC |
| P13 | PFMRPL23-5FOUT | GAGCTTTGTTCGATATAC |
| P14 | PFMRPL23-5HRFWD | CTGGTTACCTAGATATCAACCGCGGATATGAACATGAAGTAC CGGATAG |
| P15 | PFMRPL23-5HRREV | ATGTCGACCTGCAGTGAAGGTGGTTTCCACTTTGAAGAAGAT GACTTCTTGATATTTTTATTTTCACTTGAATATTCAATTGTTTC |
| P16 | PFMRPL23-3HRFWD | ATCCGCGGTCTTAAGGCTTATTTTGTACATATTAG |
| P17 | PFMRPL23-3HRREV | CTGGTTACCATCAGAATTAAATATATACACATTC |
| P18 | PFMRPL23-3FOUT | CGCACATTCTAGTTTGATATTTTG |
| P19 | PFMRPL23_gRNA1 | CATATTAAGTATATAATATTGAAAAATCGTCAAGCAGTAAAG TTTTAGAGCTAGAAATAGC |
| P20 | PFMRPL23_gRNA1_N21 | AAAAATCGTCAAGCAGTAAAG |
| P21 | PFMRPL23_gRNA2 | CATATTAAGTATATAATATTGATTTTTAATTATGTACTATAGT TTTAGAGCTAGAAATAGC |
| P22 | PFMRPL23_gRNA2_N20 | ATTTTTAATTATGTACTATA |
| P23 | N20 CHECK REV | ATATGAATTACAAATATTGCATAAAGA |
